# Functional dissection of ultraconserved non-coding regions present near the *Hoxd* complex in vertebrates

**DOI:** 10.64898/2026.10.15.759270

**Authors:** Sonu Yadav, Anil Kumar, Rakesh K Mishra

## Abstract

Ultraconserved non−coding elements (UCEs) exhibit near−perfect sequence conservation across vertebrate genomes, implying severe evolutionary constraint and essential biological functions. Near the vertebrate Hoxd complex, a master regulator of axial patterning and limb morphogenesis, three Conserved Regions (CR1, CR2, and CR3), each longer than 200 bp, show exceptional sequence conservation and drive robust regulatory activity in transient reporter assays. To investigate their endogenous in vivo functions, we used CRISPR−Cas9 genome editing to generate a series of deletions of varying sizes spanning CR2 and CR3, individually and in combination, within *hoxd* complex in zebrafish. Screening thousands of founder embryos revealed a clear size and conservation−dependent relationship between deletion extent and viability: complete deletion of core CR2, CR3, or the combined CR2/3 loci could not be transmitted through the germline, whereas the smaller and least−conserved deletion, a 626−bp subregion of CR2, was the only allele to yield a stable, homozygous−viable line. Our results indicate that the core ultraconserved sequences are developmentally indispensable, whereas flanking, less conserved sequences can be removed without consequence. Detailed phenotypic and molecular characterization of the mutant embryos revealed normal primary and secondary body axis formation, intact somite segmentation, and unperturbed craniofacial and pectoral fin development. Expression of *hoxd* complex genes showed no significant change in mutant embryos by RNA−ISH or RT−qPCR, although RT−qPCR revealed a consistent, modest upward trend at 24 and 48 hpf, raising the possibility that the deleted subregion normally contributes a weak repression of overall *hoxd* complex genes transcription. While this study reveals some functional aspects of ultraconserved elements, much more remains to be uncovered.

## Introduction

*Hox* genes are a highly conserved set of transcription factors that determine positional identity along the anterior−posterior axis and contribute to segmental identity and body plan organization in all bilaterians. *Hox* genes are expressed in a collinear manner along the body axis to determine body segment identity (Lewis, 1978; Duboule & Dollé, 1989). Any perturbation in the expression domain of *Hox* genes leads to homeotic transformation (Schneuwly et al., 1987). *Hox* complexes are subject to remarkably tight regulatory control maintained by an intricate network of *cis*− and *trans*−acting regulators, including Polycomb and Trithorax response elements (PREs/TREs), enhancers, silencers, insulators or boundary elements, and Polycomb and Trithorax groups of proteins (PcG and trxG) (Orlando & Paro, 1993; Krumlauf, 1994). Throughout evolutionary history, the genomic architecture of these clusters underwent dramatic structural reorganization (Garcia−Fernàndez, 2005). Invertebrate *Hox* complexes are generally loosely organized, showing less compaction due to the presence of intervening non−*Hox* genes, extensive repetitive elements, and divergent transcriptional orientations. In contrast, *Hox* complexes in vertebrates are more compactly organized, with no intervening genes, all transcribed in the same orientation, contain fewer introns, and have fewer repetitive elements (Darbellay et al., 2019). The compact nature arises from shorter intergenic regions in *Hox* complexes compared to those in invertebrate *Hox* complexes. Despite all this, the expression pattern remains well−organized and conserved, suggesting that strict regulatory mechanisms are at work. Vertebrates have four or more *Hox* complexes, which arose from whole−genome duplications during their evolution (Amores et al., 1998; Holland et al., 1994). Despite the compaction and increased number of complexes, the spatial and temporal expression profiles of *Hox* genes have remained largely conserved, suggesting conserved regulatory mechanisms that govern their expression. To map the genomic elements underpinning this conservation, our group focused on identifying highly constrained non− coding sequences clustered around *Hox* complexes. Interestingly, we observed three ultraconserved non−coding regions near the *Hoxd* complex, designated Conserved Region 1 (CR1), CR2, and CR3, each more than 200 bp long non−coding sequences and almost perfectly conserved across the human, mouse, and rat genomes (Sabarinadh et al., 2003). Later comparative genomic analysis across diverse vertebrate species revealed hundreds of non− coding sequences, termed Ultraconserved Elements (UCEs). These elements are predominantly non−coding, residing in intergenic or intronic regions, and are often clustered near genes involved in developmental processes (Bejerano et al., 2004). The level of conservation exhibited by UCEs is remarkably higher than that observed even in protein− coding genes, suggesting intense purifying selection pressures that tolerate no mutations. The remarkable conservation of these elements raises important questions about their biological significance: Are they mutational cold spots? Do they play vital regulatory functions in vertebrate development? Additionally, what specific role necessitates such sequence conservation? Comparing derived allele frequencies within UCEs confirms that mutation rates in these regions are typical for the genome, but fitness costs prevent mutation fixation (Katzman et al., 2007). UCEs are reported to have a high density of transcription factors binding. Multiple overlapping transcription factors binding to UCEs may impose functional constraints, thereby reducing mutation rates at UCEs (Viturawong et al., 2013). Polymorphisms in UCEs are linked to various cancers. Human genetic studies further show that polymorphisms or copy−number variations in UCE regions are heavily depleted in healthy populations but are highly prevalent in various cancers, underscoring their functional relevance to cellular homeostasis (Derti et al., 2006; McCole et al., 2014). Moreover, UCEs are frequently associated with long non−coding RNAs (lncRNAs), termed transcribed UCEs (T− UCEs), which may participate in epigenetic regulation, chromatin looping, or RNA−mediated gene silencing (Calin et al., 2007; Olivieri et al., 2016). Early studies reported that UCEs function as enhancers, silencers, or other cis−regulatory modules that fine−tune gene expression during embryogenesis (Kushawah & Mishra, 2017; Maas et al., 2011; Shin et al., 2005; Visel et al., 2007). Importantly, disruptions to UCEs, such as deletions or mutations, have been linked to developmental anomalies in model organisms and various cancers, as well as to their prognosis (Dickel et al., 2018; Pereira Zambalde et al., 2020). Beyond classic enhancer activity, recent research highlights UCEs as architectural anchors involved in higher−order chromatin looping, epigenetic modifications, and transcriptional output (Colwell et al., 2018; McCole et al., 2018).

Transient reporter assay−based studies have cataloged the regulatory potential of CRs, which act as strong repressors in cell−based and zebrafish reporter assays (Kushawah & Mishra, 2017). However, their precise *in vivo* requirements, target genes, and mechanisms of action remain unresolved. Understanding whether these elements are universally indispensable or exhibit functional redundancy requires systematic, stable genetic ablation within an endogenous, living system. In this study, we addressed these questions through a dual bioinformatic and genetic approach in zebrafish. We first leveraged public datasets (UCSC Genome Browser, ChIP−seq, and ChromHMM) to dissect the epigenetic and transcriptional signatures of the CR loci across multiple vertebrate species. Our in−silico analyses revealed that CR2 and CR3 act as bivalently poised regulatory hubs, predominantly occupied by Polycomb Repressive Complex (PcG) proteins (such as SUZ12 and EZH2) in non−expressing tissues, while acquiring active histone marks (H3K4me1/3) in developing structures like the embryonic limb. Driven by these findings, we deployed CRISPR−Cas9 genome editing to systematically delete the CR2 and CR3 loci, both individually and in combination, to assess their essentiality during *in vivo* embryonic development. We report that core ultraconserved sequences within these loci are not fully dispensable, indicating strong selective pressures against their loss during early development. We report that no heritable allele removing the ultraconserved core of either locus could be recovered, an outcome consistent with strong selective pressure against their loss during early development. However, we successfully established and analyzed a stable, homozygous zebrafish line carrying a 626−bp deletion in a less−conserved subregion of CR2 (termed CR2Δ626). By tracking embryogenesis, somitogenesis, craniofacial skeletal development, and pectoral fin morphogenesis in this line, alongside qualitative and quantitative spatial−temporal expression profiling of the *hoxd* complex, this work provides new insights into the functionality of CR, regulatory redundancy, and also emerging transcriptional properties of vertebrate ultraconserved elements.

## Results

### Bioinformatic analysis of CRs loci

We performed a comprehensive analysis of CR1, CR2, and CR3, utilizing publicly available data from the UCSC Genome Browser, to investigate their sequence conservation, transcriptional potential, regulatory interactions, and epigenetic landscapes across various species. We performed a multiple sequence alignment of the region containing CR1, CR2, and CR3 from several representative vertebrate species. We found that while extended CR1, CR2, and CR3 are highly conserved across all tested species, the core CRs (the green transparent box in Figure 1) exhibit extraordinary sequence conservation (Supplementary Table 1 and Figure 1A).

**Figure 1:**
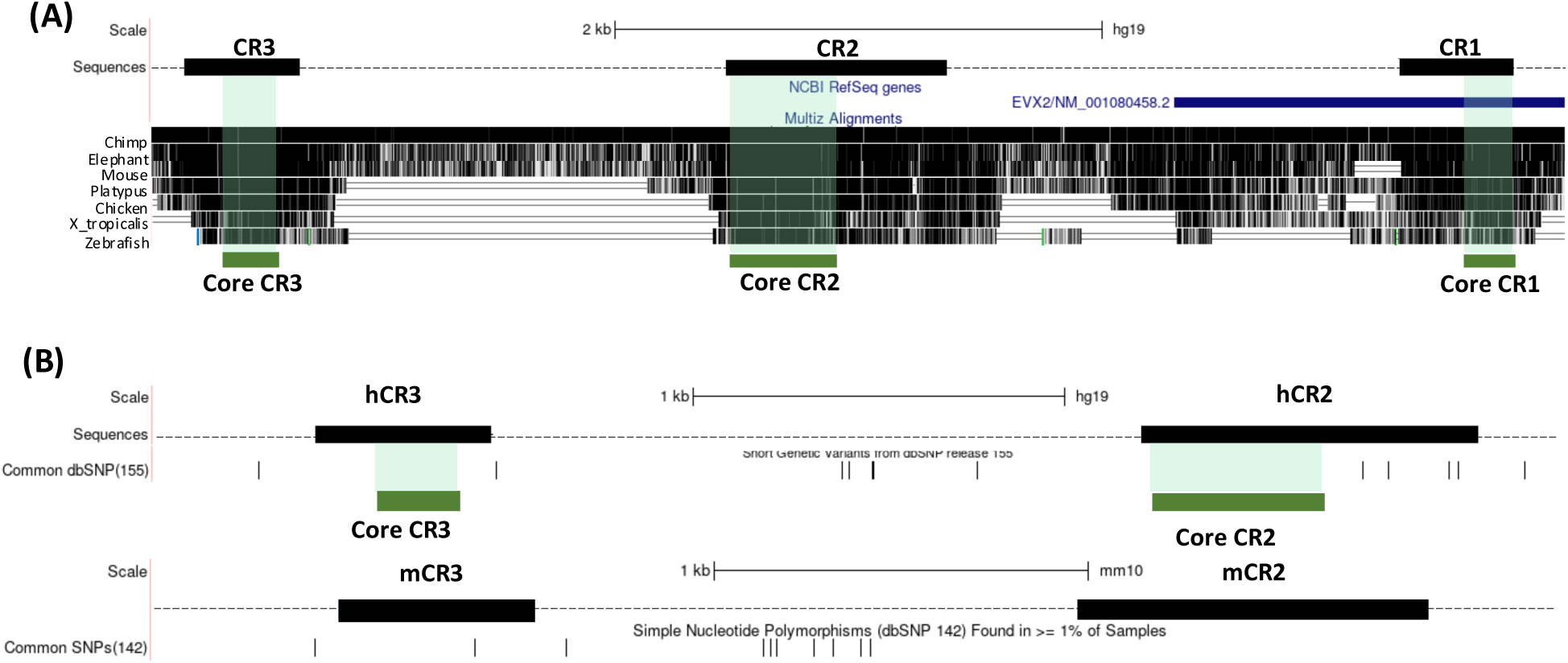
Evolutionary conservation and genetic variations in CR loci. (A) Multiple sequence alignment of CRs in vertebrate species. The top track, labelled “sequences,” shows the extended CR sequences (black boxes). Below the CRs is the RefSeq genes track. The bottom track displays multiple sequence alignments of CR loci across vertebrate species. The density of black bars in the alignment track corresponds to the level of sequence conservation in the alignments. The green transparent box over the alignment track and the green solid box beneath indicate the core CRs. (B) Sequence polymorphism in CRs loci. The short genetic variants from dbSNP release 155 are mapped below the “sequences” track. Common SNPs, variants with at least 1% minor allele frequency, are represented by vertical bars. Common SNPs mapped to the mouse CR locus from dbSNP build 142.

In higher vertebrates, the extended conservation of sequences on either side of core CRs forms an inverted pyramid of conservation from lower to higher vertebrates. This indicates that very few mutations were tolerated in core CRs during evolution, with increasing conservation pressure on the flanking sequences as we move to higher vertebrates. This observation suggests the accumulation of additional functional information in the DNA sequences surrounding core CRs in advanced vertebrates. From here onwards, we use the term “CR” for extended CR unless described otherwise. We further figured that CR1 is less conserved than the other two CRs and is located within the 3’ UTR of the *Evx2* gene (Figure 1A). In the current study, we aim to understand the function of non−coding CRs that do not overlap with any genes. Therefore, we focused on CR2 and CR3. Earlier studies have reported that UCEs are depleted of commonly found genetic variants. Therefore, we analyzed the presence of common SNPs over CR loci. We observed multiple SNPs outside CRs (Figure 1B). CR3 and core CR2 do not contain any SNPs, whereas the remaining part of the CR2 has a few SNPs. Similarly, in the mouse common SNPs dataset, we observed only one SNP in CR3, and the remaining SNPs were located outside CRs (Lower track in Figure 1B). We examined the transcriptional activity of CR2 and CR3. We identified expressed sequence tags (ESTs) from CR3 in 8−cell−stage mouse embryos, stage 36 of chicken embryos, and Xenopus tadpoles (Supplementary Figure 1).

UCEs are reported as binding hubs for transcription factors. Genomic mapping in human and mouse cell lines and tissues demonstrated high−density transcription factor−binding peaks clustered over CR2 and CR3 (Supplementary Figures 2A and 2C). The vast majority of these binding factors are components of the Polycomb Group (PcG) repressive complex, including SUZ12, EZH2, RNF2, CBX8, and BMI1, which establish repressed chromatin states (Supplementary Figure 2B). Additionally, the chromatin remodeler SMARCA4 and the developmental transcription factor TCF2 (HNF1B) occupy specific subregions of CR2 (Supplementary Figure 2B).

**Figure 2:**
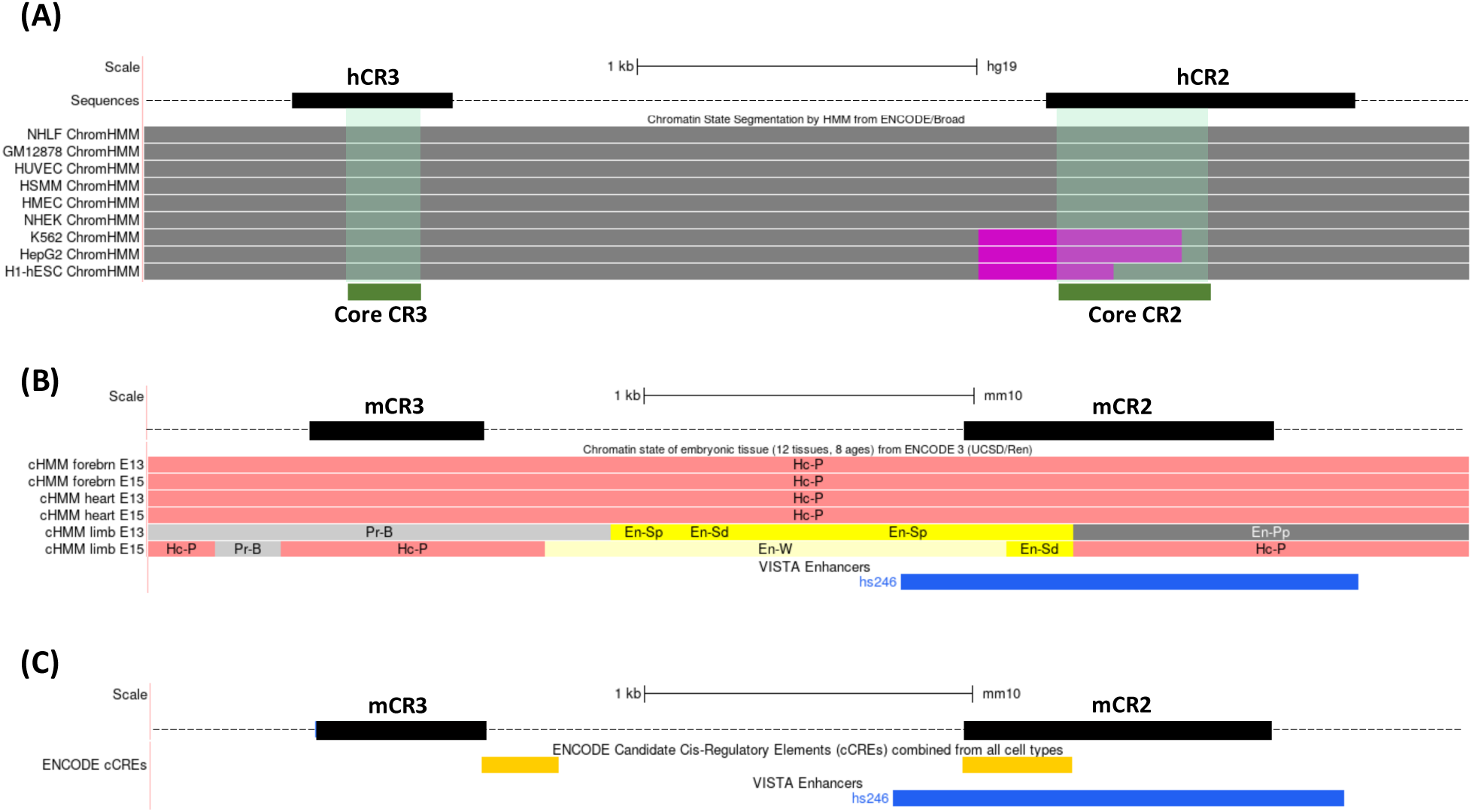
Regulatory landscape of CRs loci. (A) Regulatory functionality predicted by integrating ChIP seq data for CTCF and histone modifications (H3K4me1/2/3, H3K9ac, H2K27ac, H2K9me3, H3K27me3 and H3K36me3) using Hidden Markov Model (HMM) in different human cell types. Grey-Polycomb repressed and purple-Inactive/poised promoter (B) Regulatory functionality predicted by integrating ChIP-seq data of histone modifications (H3K4me1/2/3, H3K9ac, H2K27ac, H2K9me3, H3K27me3 and H3K36me3) using HMM in mouse embryonic tissues. These states were combined and given different colours to represent different functional elements. Hc-P: heterochromatin Polycomb associated, En-Sp: Enhancer, Strong TSS-proximal, En-Sd Enhancer: Strong TSS-distal, Pr-B: Promoter, Bivalent, En-Pp: Enhancer, Poised TSS-proximal (C) Candidate Cis-Regulatory Elements (cCREs) identified based on DNase hypersensitivity, histone modification (H3K4me3 and H3K27Ac) and CTCF binding sites across samples analysed in ENCODE project. VISTA enhancers (Blue bars) comprise experimentally validated enhancers in transgenic mice.

Histone modification profiles and ChromHMM models indicated that CR regions are predominantly repressed via H3K27me3 methylation and EZH2 recruitment across most human cell lines (Supplementary Figure 3A) and mouse tissues (heart tissue is shown as an example in Supplementary Figure 3B). However, specific cell types (K562, HepG2, and H1− hESC) exhibit a poised promoter signature overlapping with core CR2, characterized by the simultaneous presence of repressive H3K27me3 and active marks (H3K4me1/2/3 and weak H3K27ac) (Figure 2A). In embryonic mouse limb tissues (E13 and E15), the chromatin architecture shifts dramatically: activating H3K4me1/2/3 marks enrich across the loci while repressive H3K27me3 marks clear away (Supplementary Figure 3B). Under these conditions, ChromHMM predicts distinct enhancer−like activity for CR2 and a bivalent promoter signature for CR3 (Figure 2B). This distal enhancer prediction is supported by the ENCODE candidate Cis− Regulatory Elements (cCRE) study and validated by *in vivo* mouse reporter assays, which show that CR2 (hs246) drives tissue−specific expression in the embryonic forebrain, hindbrain, and neural tube (Figure 2C). In summary, our analyses indicate that CRs are highly constrained vertebrate elements with dynamic, tissue−specific regulatory potential.

### CRISPR-Cas9-mediated deletion of CRs in zebrafish

Most ultraconserved elements are located near the developmental genes. In a mouse transgenic reporter assay, a major fraction of UCEs was reported to have enhancer−like activity, driving reporter gene expression in a manner similar to that of nearby genes. Reporter−assay− based studies in human cell lines and zebrafish embryos revealed that CRs also play a regulatory role. But their target genes and the mechanism of action were not revealed. To further understand the *in vivo* function of CR2 and CR3, we planned to delete the entire locus spanning CR2 and CR3, the entire CR2 locus, the region between CR2 and CR3, CR2 and CR3 individually, and smaller deletions in CR2 and CR3. Based on the presence of the PAM sequence NGG, with high efficiency scores and minimal off−targets, we designed multiple gRNAs (Figure 3A) and used them in combination to target various regions (Supplementary Table 2). From this large−scale screen of thousands of founder embryos, we identified founder fish with deletions: 1899 bp spanning the entire CR2, with a region between CR2 and CR3 (purified deletion band of size 300bp in Figure 3B); 1111 bp covering the entire CR2, with some flanking regions (Figure 3C); and a smaller deletion of 626 bp in CR2 (Figure 3D). The deletions were confirmed by PCR−based genotyping and subsequently by Sanger sequencing of the deletion band. We further screened at least 800 F1 progenies from the respective deletion fish lines but obtained germ−line transmission of the deletion of 626bp in CR2 and subsequently maintained a homozygous deletion line, named CR2Δ626. We did not obtain any founder fish with CR3 deletion. The deletion targets and the screening results are summarised in Figure 3E. We further examined the sequence conservation of the deleted 626bp of CR2 across vertebrate species and found this subregion to be comparatively less conserved that the rest of CR2 (discussed in a later section).

**Figure 3:**
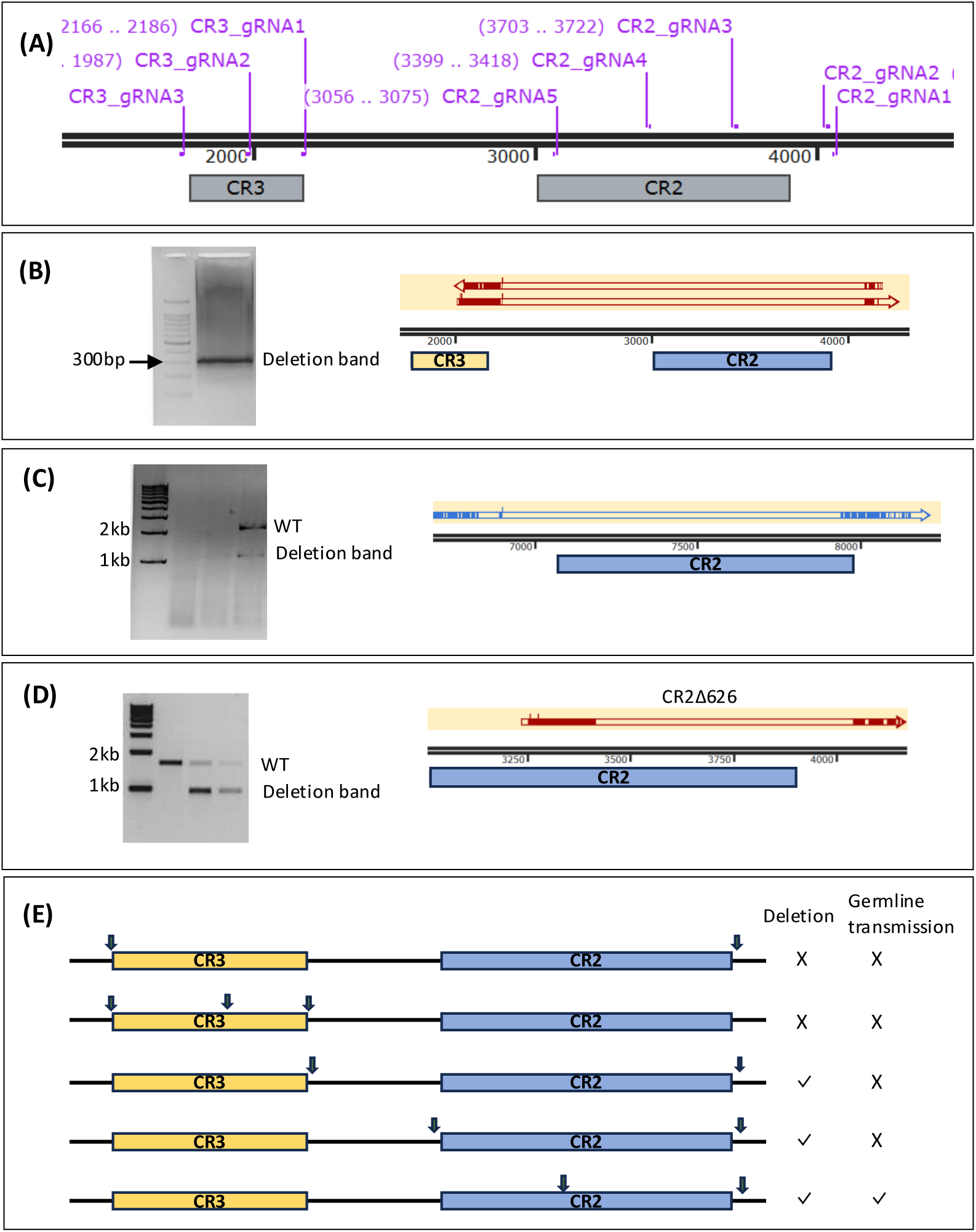
Deletion of CRs using the CRISPR-Cas9 technique. (A) Guide RNA targets for CR2 and CR3 (B) Founder fish having deletion of CR2 and the region between CR2 and CR3. The agarose gel on the left shows the purified deletion band of approximately 300 bp in size, and the right side displays the alignment of the deletion band sequence with the wild-type sequence in SnapGene. (C) Screening of founder fish containing the whole CR2 deletion of around 1kb in size and sequence confirmation of the deletion band. (D) Screening of progenies from parents with a partial deletion of CR2, approximately 600 bp in size, and sequence confirmation of the deletion band. (E) Summary of CRISPR deletion of CRs and their germ-line transmission

Taken together, this deletion series revealed a relationship between sequence conservation and transmission success: germline transmission was obtained only for the least conserved deletion tested (626 bp), whereas deletions encompassing core CR2 and/or CR3 sequences were never transmitted despite extensive founder screening (Figure 3E, Supplementary Table 2). This pattern points to selective pressure acting specifically against loss of the core ultraconserved sequence. Although we cannot rule out inefficient cutting at the target sites by the gRNAs, and the limited founder recoveries.

### Role of deleted CR2 in early development of zebrafish

Although we were unable to obtain a complete CR2 deletion, we successfully deleted 626 bp of CR2, which was also germ−line transmitted to the next generation and maintained as a homozygous deletion line. All the experiments were performed in the homozygous deletion line. We used the term “CR2Δ626” to refer to the partially deleted CR2. Since CR2 is near the *hoxd* complex, we hypothesized that it may regulate *hoxd* genes expression. *Hox* genes are known to regulate body segmentation and body axis formation. Any misregulation may result in visible phenotypes during early development. We performed phenotypic screening in CR2Δ626 embryos but observed no developmental defects (Figure 4A). We also assessed the number and morphology of somites by staining the 2dpf embryos with the *xirp2a* gene, which marks the somite boundaries (Figure 4B). However, we did not observe any morphological differences in the somites of CR2Δ626 embryos compared with WT embryos. Also, the average number of somites was the same in both WT and CR2Δ626 embryos (Figure 4C). We also examined the Mendelian ratio to assess the survival of heterozygous and homozygous embryos (2dpf) and larvae (15dpf) and observed no significant deviation from the expected ratios (Supplementary Table 3). Further, there was no significant difference in the viability, breeding rate, and clutch size of the homozygous line as compared to WT. These embryos further developed into healthy, fertile fish with no visible phenotypic defects. Based on the above information we can say that the deleted subregion of CR2 is not required for segmentation, body−axis formation, and survival, at least under the tested conditions.

**Figure 4:**
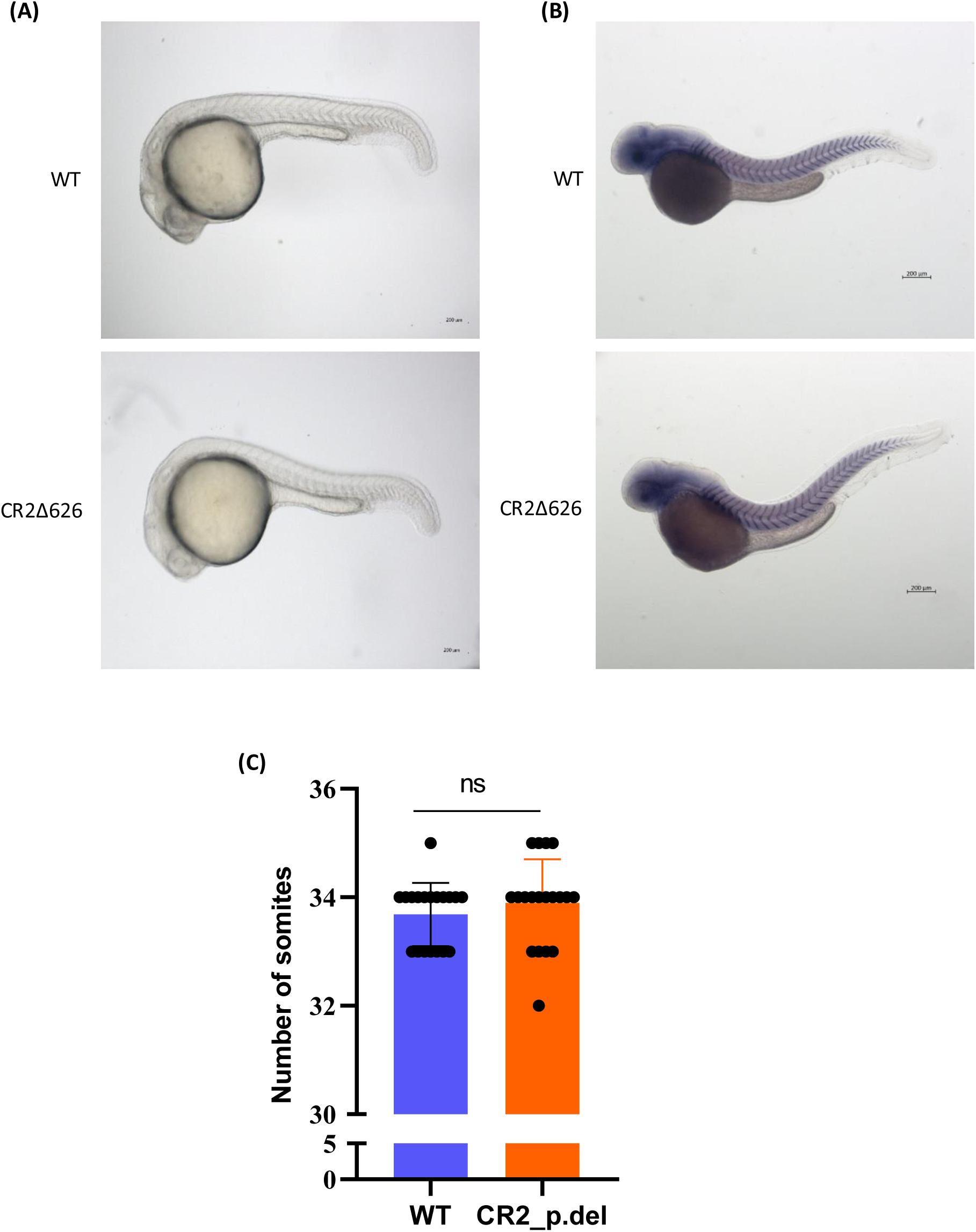
Body axis and segmentation in CR2Δ626 embryos. (A) Morphology of 24hpf wild and CR2Δ626 embryos. B) Somites in 48hpf wild type and CR2Δ626 embryos where somite boundaries are stained by xirp2a. C) Somite count in 48hpf wild type and CR2Δ626 embryos, n=19 embryos per genotype.

### Role of deleted CR2 in cranio-facial features and pectoral fin development

Apart from their role in body axis and segment determination, *Hox* genes are involved in the development of pharyngeal arches and craniofacial features (Minoux & Rijli, 2010; Trainor & Krumlauf, 2001). Misregulation of *Hox* genes may lead to defects in the craniofacial bone and cartilage. Thus, to study the role of CR2 in bone and cartilage development, we stained 10 dpf old WT and CR2Δ626 fish with alizarin red and alcian blue stains that are known to stain bone and cartilage, respectively. We observed properly stained bones and cartilage, but found no distinct deviation in the CR2Δ626 compared with WT fish (Figure 5A). In vertebrates, *Hox* genes not only regulate the development of the primary axis but are also required for the development of the secondary axis (limbs). In vertebrates, expression of *Hox* genes follows spatial−temporal collinearity during limb development, a feature not observed in invertebrates. CRs are located near the *hoxd* complex, which is known to be crucial for limb development. Thus, we hypothesized that CRs might be important for regulating *hoxd* genes during limb development, and that their deletion might affect limb development. We investigated whether the deleted subregion of CR2 has any function in the pectoral fin development of zebrafish. To study the development of the pectoral fin, we performed alcian blue staining to mark the cartilage of the pectoral fin of 10dpf WT and CR2Δ626 fish. CR2Δ626 fish showed no morphological defects in the pectoral fins as compared to WT (Figure 5B). Our findings suggest that the deleted subregion of CR2 is not required for pectoral fin development.

**Figure 5:**
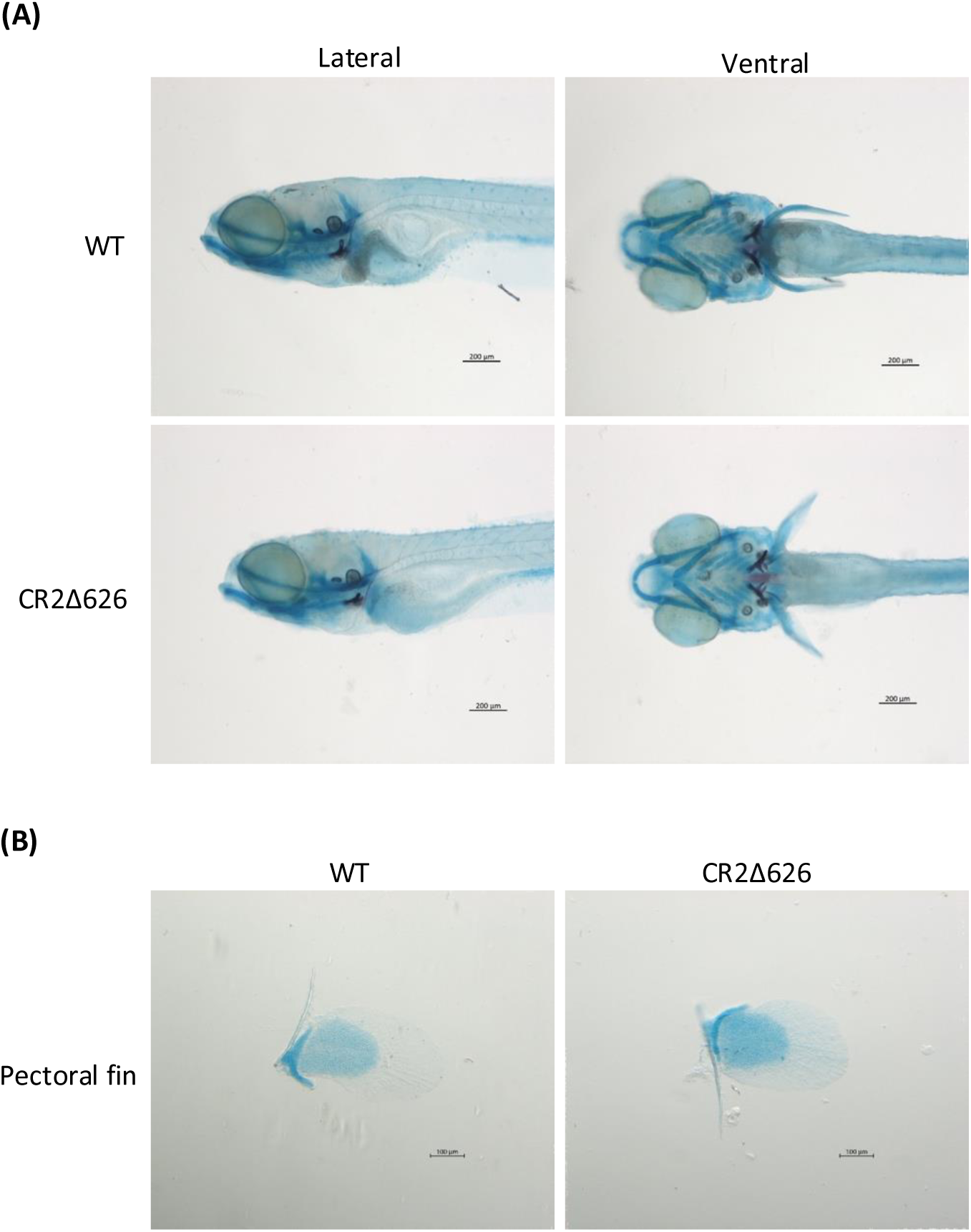
Craniofacial features and pectoral fin development. (A) Cranio-facial bone and cartilage stained by alizarin red and alcian blue, respectively, in 10 dpf WT and mutant fishes (B) Pectoral fins of 10dpf old WT and CR2Δ626 mutant fishes stained with alcian blue. Dark blue colour is the endoskeletal disc and the transparent tissue surrounding it is the fin fold.

### Expression of *hoxd* complex genes in partially deleted CR2 embryos

Studies have shown that the centromeric and telomeric flanks of the *Hoxd* complex contain regulatory regions that control the spatiotemporal expression of *Hoxd* genes (Montavon et al., 2011; Rodríguez−Carballo et al., 2017). CRs are adjacent to *Hoxd* complexes and have regulatory activity, suggesting they may regulate hoxd gene expression. In 24hpf embryos, *hoxd* genes are well established in the primary axis. In WT embryos, the *hoxd3* gene is expressed in the hindbrain region with weaker expression towards the posterior end. The *hoxd4* gene is expressed in the hindbrain rhombomeres, with a sharp anterior boundary. Expression of both these anterior *hoxd* genes remains unchanged in CR2Δ626 embryos (Figure 6). In WT embryos, *hoxd9* and *hoxd10* genes are expressed in the middle of the primary body axis, and the expression remained unchanged in CR2Δ626 embryos (Figure 6). The posterior genes *hoxd11*, *hoxd12*, and *hoxd13* are expressed posteriorly in the tail region, thereby maintaining the expression collinearity along the body axis. Additionally, *hoxd13* is also expressed in the cloacal aperture (marked by red arrows in Figure 6). The expression pattern of these genes in CR2Δ626 embryos remained the same as in WT. The expression of the *evx2* gene also remains unchanged along the body axis (Figure 6). We further studied the expression profiles of *hoxd* genes at 48 hpf in the primary and secondary axis (pectoral fin buds). The expression pattern of *hoxd* genes in the primary axis of 48hpf embryos remained the same as mentioned above upon CR2 deletion (Supplementary Figure 4). In WT pectoral fin buds, *hoxd* genes followed spatial collinearity, where *hoxd9* and *hoxd10* are expressed in the anterior region of the pectoral fin bud (indicated by red arrows in Figure 7), whereas *hoxd11*, *hoxd12*, and *hoxd13* are expressed in posterior regions of the fin bud (indicated by yellow arrows in Figure 7). CRs might play a role in hox gene regulation at earlier stages, so we also assess their expression in 16hpf embryos. But we do not find any significant difference in the expression domain of CR2Δ626 embryos compared to WT (Supplementary Figure 5). However, changes in expression levels cannot be quantified by RNA−ISH unless they are substantial. Therefore, we performed RT−qPCR to assess relative expression in WT vs. CR2Δ626 embryos at 16 hpf, 24 hpf and 48 hpf, using four independent embryo pools per genotype at each stage. We did not find any significant difference in *hoxd* complex gene expression levels at 16 hpf (Supplementary Figure 6A); however, we observed modest increases in the expression of all *hoxd* genes at 24 hpf and 48 hpf (Supplementary Figure 6B and 6C), but the changes were not significant (Supplementary Table 4). Overall, we conclude that the deleted subregion of CR2 does not significantly affect *hoxd* complex gene expression during early development, although the consistent, modest upward trend observed at 24 and 48 hpf is intriguing and is considered further in the Discussion.

**Figure 6:**
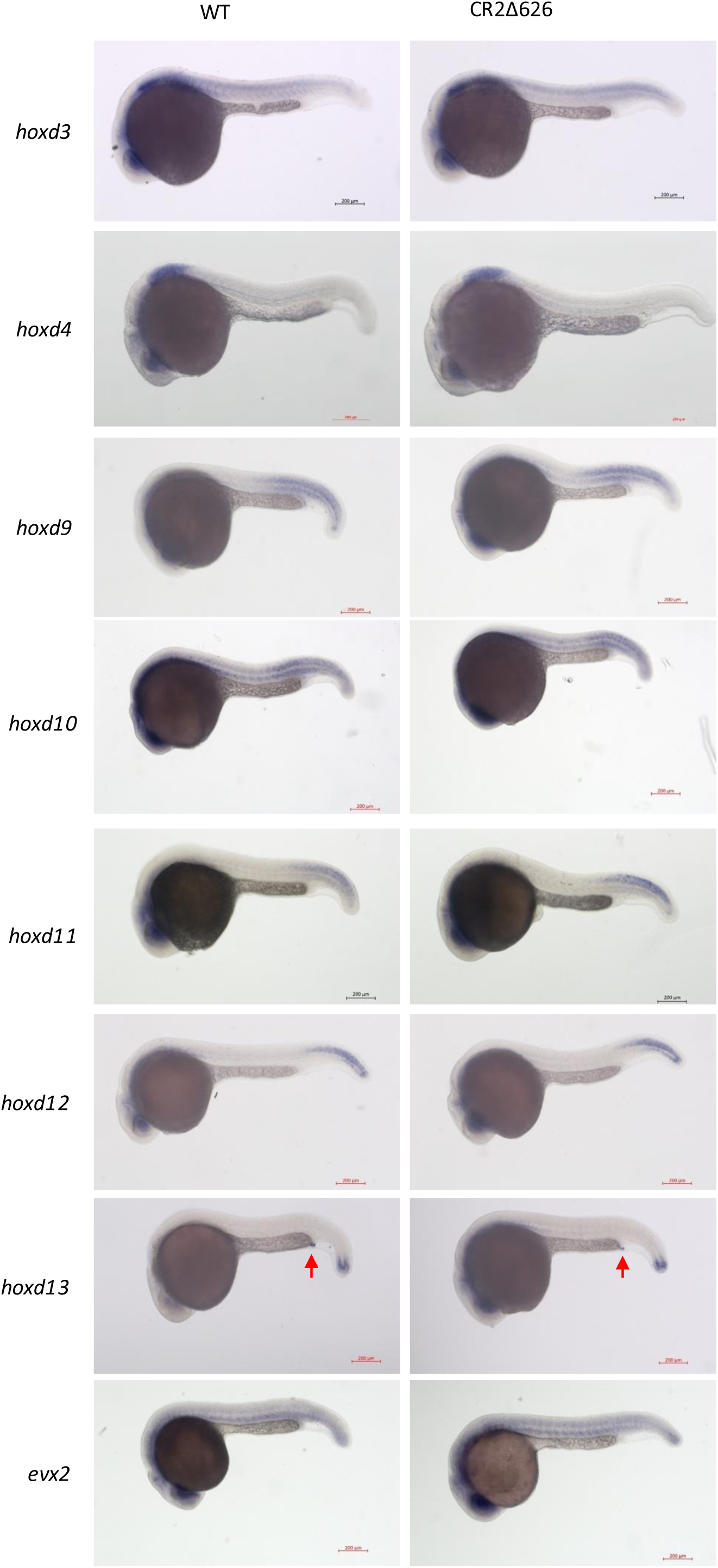
Expression of *hoxd* complex genes in 24 hpf CR2Δ626 embryos. RNA-ISH of all the *hoxd* genes and *evx2* gene in 24 hpf WT and CR2Δ626 embryos. Red arrow represents the cloacal aperture.

**Figure 7:**
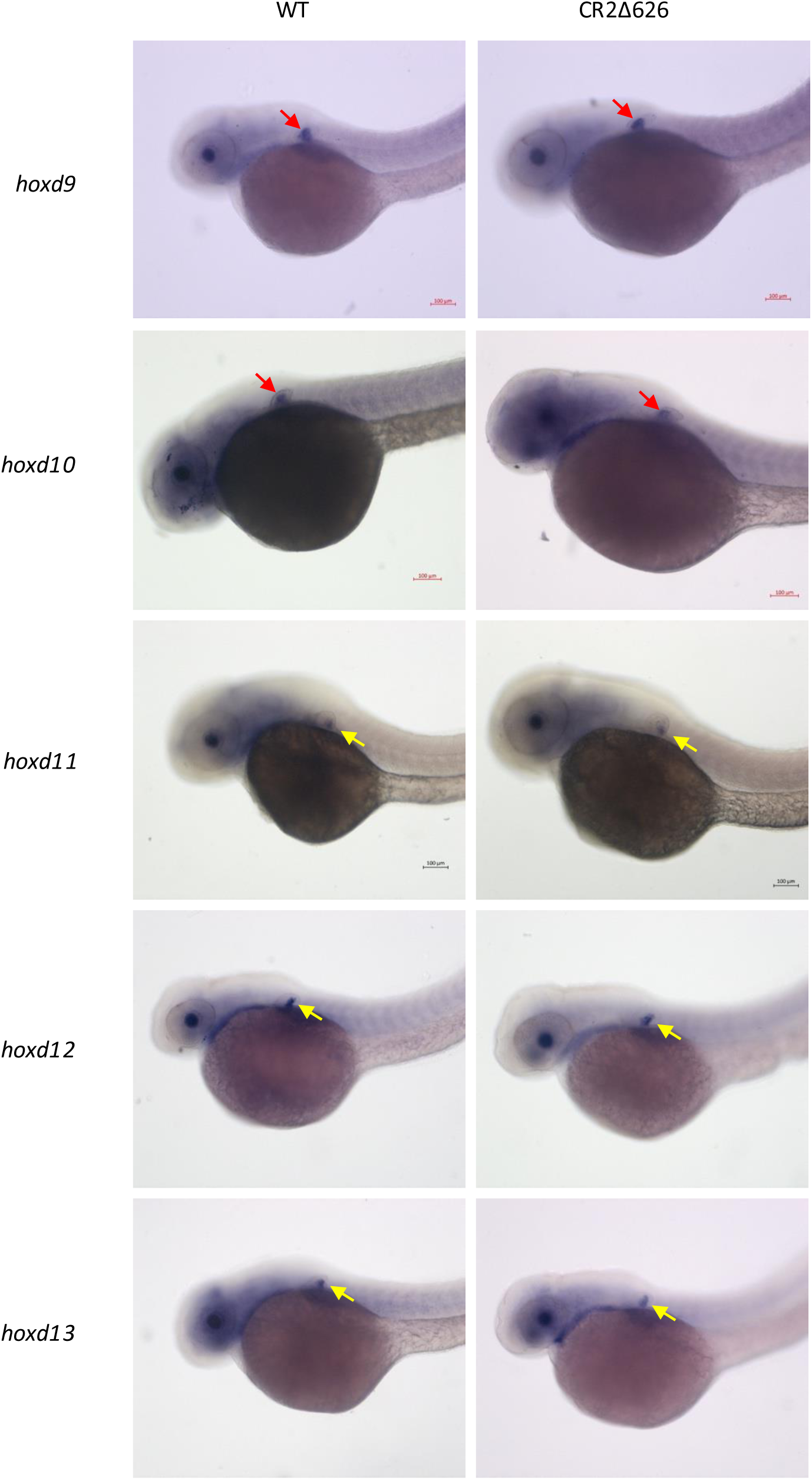
Expression of *hoxd* genes in CR2Δ626 embryos. RNA-ISH of *hoxd* genes gene in 48hpf wild type and CR2Δ626 embryos. Red arrows indicate *hoxd* genes expression in the anterior pectoral fin bud and yellow arrows indicate *hoxd* genes expression in the posterior pectoral fin bud.

### Analysis of the deleted subregion of CR2

CR2Δ626 embryos showed no morphological defects and no detectable change in the spatial expression of *hoxd* complex genes, with only a modest, non−significant increase in hoxd complex expression levels at 24 and 48 hpf. To further investigate the deleted 626−bp subregion from the rest of CR2, we returned to the comparative and epigenomic datasets described above (Figure 8). In the Multiz alignments, core CR2 is near−perfectly aligned across the full span of species surveyed, from chimp through to *X. tropicalis* and zebrafish. Alignment over the deleted subregion is comparably strong in chimp, elephant, mouse and platypus, becomes patchy in chicken, and is largely absent in *X. tropicalis* and zebrafish. Conservation across the extended element is therefore not uniform: it is vertebrate−wide at the core and progressively lineage−restricted toward the flank. Population−level and factor−binding data follow the same asymmetry. Common variants from dbSNP release 155 map to the deleted subregion while the core−containing remainder is devoid of them, indicating that the core has been under stronger purifying selection. ReMap density, which aggregates transcription factor binding peaks across datasets, peaks sharply over core CR2 and declines across the deleted subregion. The two parts also differ in chromatin behavior. Across the five cell lines segmented by ChromHMM, a poised/inactive promoter−like signature is called over core CR2 in K562, HepG2 and H1−hESC, whereas HUVEC and GM12878 remain repressed throughout. The deleted subregion carries no such signature on any of the five lines, remaining Polycomb− repressed throughout. The deleted subregion is thus the less constrained and less factor− bound of the two, yet the more uniformly repressed.

**Figure 8:**
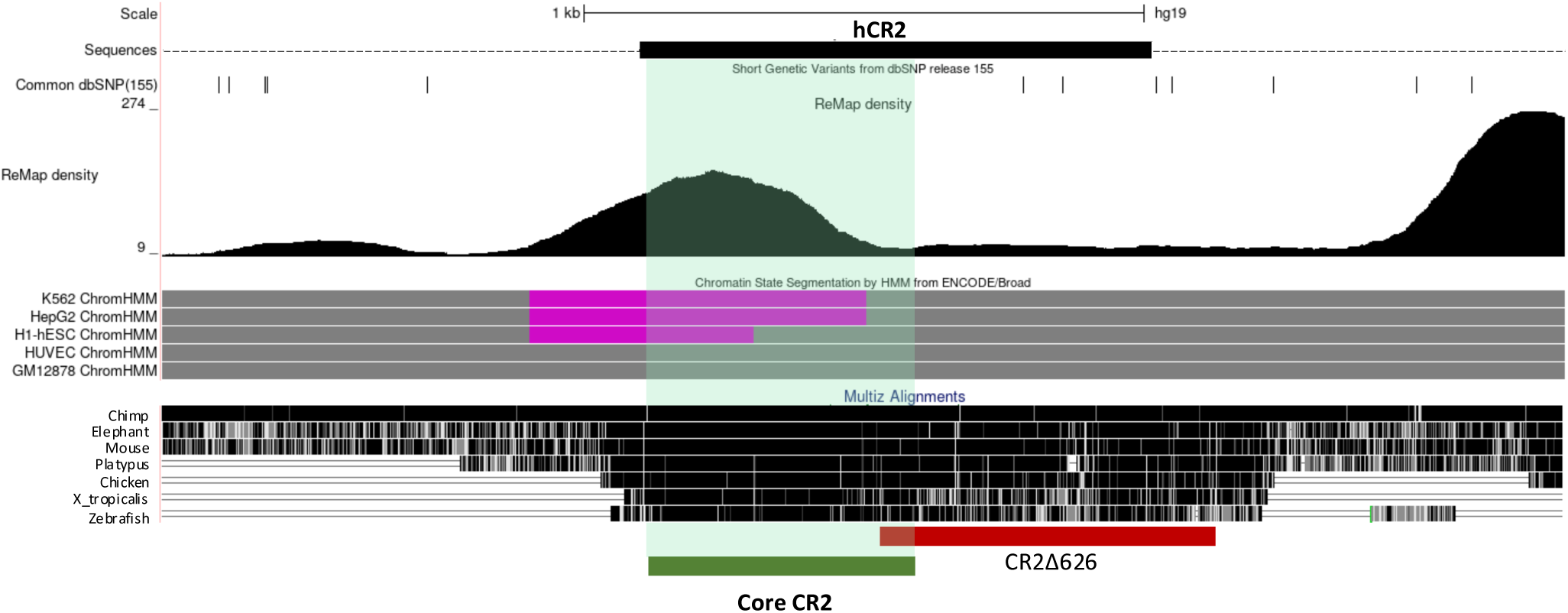
Comparative and epigenomic features of the extended CR2 locus. UCSC Genome Browser view of the extended CR2 region (hg19). Tracks, from top: scale bar; extended CR2 sequence; common short genetic variants from dbSNP release 155, shown as vertical bars; ReMap density of aggregated transcription factor binding peaks (scale 9–274); chromatin state segmentation by HMM (ENCODE/Broad) in K562, HepG2, H1-hESC, HUVEC and GM12878 cells, with grey denoting Polycomb-repressed states and purple denoting poised promoter; and Multiz alignments across mentioned species. The green bar and green transparent box mark core CR2; the red bar marks the subregion deleted in the CR2Δ626 line.

Considered alongside the failure of core CR2, CR3 and combined CR2/3 deletions to be transmitted through the germline, these observations indicate that functional constraint within the extended CR2 is concentrated in the core, and that the flanking subregion removed in CR2Δ626 is dispensable for zebrafish development, although its invariant repressive chromatin signature suggests it is not simply inert.

### Transcriptional status of CRs

A large number of UCEs are known to be transcribed into non−coding RNA (T−UCEs). Genetic variations in these T−UCEs are also associated with various cancers and other diseases. We observed ESTs from CR3 in mouse, chicken, and xenopus samples. Since CRs are conserved across vertebrates, their transcriptional status may also be conserved across vertebrates. Therefore, we aimed to investigate the transcriptional status of CRs (CR2 and CR3) in zebrafish embryos. We performed RNA−ISH for CR2 and CR3 during early development using both sense and antisense probes, prepared from CR2 and CR3 DNA templates, respectively (Figure 9A). We obtained a strand−specific signal in both CR2 and CR3 probes, suggesting transcription direction opposite to evx2 and the same as *hoxd* genes transcription direction. RNA from both CR2 and CR3 exhibited similar expression profiles, suggesting that a single RNA transcript may be transcribed through both CR2 and CR3. But it needs further investigation. According to the signal, CR RNAs were observed as early as the 64−cell stage, suggesting maternal deposition (Figure 9B and C). Both CR RNAs were expressed throughout the embryo until gastrulation began. Towards the onset of gastrulation, RNA signal started to concentrate more around the shield and gradually towards the anterior region of the embryo. At the 24 hpf stage, CR RNAs were expressed in the eye, brain, and anterior few somites (Figure 9B and C). The sense strand does not show much signal in most embryos in the tested samples (24hpf embryos shown in Figure 9D as a representative example). To summarize, we found strand−specific transcription of CRs in the same direction as *hoxd* genes. The expression was initially global, but became specific to the anterior region following gastrulation.

**Figure 9:**
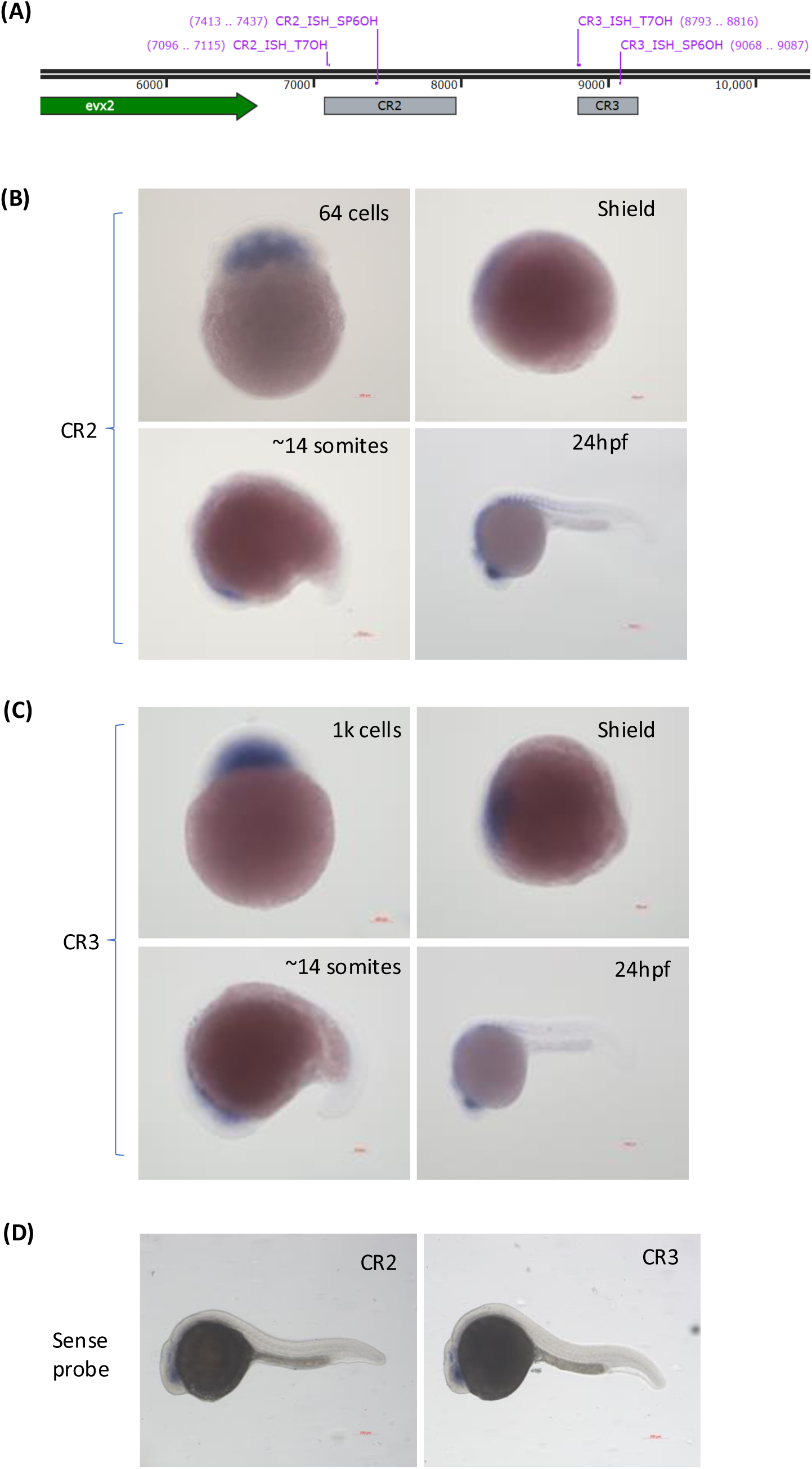
CR RNA expression profile in the early embryos. (A) Schematic of primers used to design RNA probes for CR2 and CR3 RNA-ISH. (B) CR2 RNA expression in early embryonic stages. (C) CR3 RNA expression in early embryonic stages. (D) 24 hpf WT embryos stained with labeled sense probe of CR2 and CR3.

## Discussion

In vertebrate genomes, hundreds of ultraconserved non−coding elements (UCEs) occupy highly constrained loci, yet their endogenous functions remain largely enigmatic. Near the vertebrate *hoxd* complex, three such regions — CR1, CR2 and CR3 — exhibit near−perfect sequence conservation across diverse species. Our bioinformatic analysis indicates that these loci possess context−dependent epigenetic regulatory roles: the CR loci are repressed across most human and mouse cell types, while core CR2 overlaps bivalent or poised promoter signatures in a few human cell lines and displays enhancer−like active histone marks in mouse embryonic limbs. To test their function in vivo, we targeted CRs with deletions of varying size. Mosaic founders carrying a deletion of the entire CR2, or a larger deletion spanning CR2 to CR3, never achieved germ−line transmission, and complete deletion of CR3 could not be recovered even in mosaic form. We are cautious in interpreting this, as there can be technical explanations that our data cannot exclude: inefficient cutting at the larger deletion targets, low germ−line mosaicism, and the limited number of founders recovered during genotyping. Even allowing for these technical possibilities, the repeated failure to recover a transmissible deletion of either region across independent targeting attempts strongly suggests that core CR2 and CR3 being developmentally indispensable and subject to purifying selection that tolerates no major disruption, consistent with population−level depletion of UCEs within copy− number variants and segmental duplications (Derti et al., 2006).

In contrast, the smaller CR2Δ626 allele yielded a stable line with no morphological defects in somite boundaries, craniofacial structures or pectoral fin development, and no discernible change in the spatial expression domains of *hoxd* complex genes at 24 and 48 hpf, including in the fin buds. RNA−ISH, however, is poorly suited to detecting subtle quantitative shifts, and RT−qPCR accordingly revealed a modest, statistically non−significant upward trend in *hoxd* expression at both stages. The chromatin behaviour of the two halves of the extended CR2 offers a possible explanation for its direction (Figure 8). The deleted subregion remains Polycomb−associated irrespective of cellular context, whereas the retained core alternates between repressive and poised, promoter−like states. One plausible interpretation is therefore that the deleted subregion normally contributes a weak but constitutive repressive input to *hoxd* transcription, and that its loss partially relieves this repression without crossing the threshold required for an overt phenotype, with residual silencing from the intact core CR2 and CR3 buffering the effect. Under this view, the phenotypic normality of CR2Δ626 would reflect not the functional irrelevance of the deleted subregion, but a quantitative rather than qualitative contribution to *hoxd* regulation, a distinction that assays powered to detect gross morphological or strong transcriptional change, such as ours, are not designed to resolve.

This pattern, an ultraconserved sequence whose deletion leaves the animal grossly normal, is the central paradox of the UCE field. Ahituv et al. (2007) removed four UCEs from the mouse genome and recovered viable, fertile animals without overt phenotypes, a result widely read as evidence that extreme conservation need not imply individual indispensability (Ahituv et al., 2007). That interpretation was substantially revised by Dickel et al. (2018), who showed that deletions of ultraconserved enhancers do produce phenotypes when assayed with sufficient sensitivity, but that these are frequently quantitative, tissue−restricted or expressed only in particular contexts rather than appearing as gross morphological defects (Dickel et al., 2018). In our investigation, complete removal of core CR2 or CR3 was never transmitted, placing these sequences firmly in the indispensable category and arguing against a general conclusion that ultraconservation is phenotypically inert. The viable CR2Δ626 allele, by contrast, resembles the Ahituv outcome superficially, with no gross phenotype, while the accompanying quantitative trend is more consistent with the Dickel interpretation, in which effects exist but fall below the detection threshold of standard assays. The distinction our data adds is positional: within a single ultraconserved region, the core and its extended flank behave differently, such that whether a UCE deletion appears consequential depends on how much of the core is removed. Reconciling the apparently conflicting UCE−knockout literature may therefore require attention not only to assay sensitivity but to which part of an ultraconserved region a given deletion actually removes.

Revisiting our bioinformatic data provided crucial context for why this partial deletion line was viable and phenotypically normal. Common SNPs were confined to the deleted subregion of CR2, whereas the remaining core region of CR2 and CR3 completely lacked SNPs and harbored a significantly higher density of transcription factor binding sites. Evolutionary analysis further revealed an “inverted pyramid” of conservation: while core CRs are conserved across all vertebrates, the extended flanking regions, which include the 626−bp subregion deleted in CR2Δ62, accumulated conservation primarily in higher vertebrates. We consider three, non− mutually exclusive explanations for the absence of an overt phenotype in CR2Δ626zebrafish; none is directly tested by our current data, and we flag them as hypotheses rather than conclusions. First, because this subregion is less conserved in teleosts such as zebrafish, it may lack critical functional relevance in this lineage; its greater constraint in mammals could instead reflect the acquisition of essential regulatory roles that drive mammalian−specific anatomical and physiological novelties. Throughout tetrapod and mammalian evolution, Hox clusters were repeatedly co−opted to direct specialized developmental programs, such as *Hoxc13* in hair follicle morphogenesis (Godwin & Capecchi, 1998), *Hoxa10/11* in uterine receptivity and endometrial remodeling (Taylor et al., 1998), and *Hox10* group in abdominal axial regionalization via lumbar rib suppression (Wellik & Capecchi, 2003). Crucially, while we observed no defects in zebrafish pectoral fin buds or *hoxd* expression in pectoral fins, 5’ *HoxD* regulatory landscapes in mammals are heavily deployed during Phase III late−stage limb patterning to specify the autopod (digits), a structural innovation absent in teleost fins (Montavon et al., 2011). The lack of a detectable phenotype in our teleost knockout model is therefore consistent with a model in which this subregion became functionally constrained only after its recruitment into higher−vertebrate− or mammalian−specific regulatory networks. Second, and independent of any mammalian−specific recruitment, the absence of a severe phenotype following the loss of this subregion may instead reflect functional redundancy, potentially buffered by compensatory regulatory activity from the intact CR3 locus or the neighboring core CR2 unit, as well as by broader genetic buffering networks that safeguard fundamental body plans. Because CRs are exclusive to vertebrate *hox* complexes and absent in invertebrates, they may not be strictly required for universal *Hox* functions, but rather evolved to fine−tune vertebrate−specific processes, such as maintaining secondary−axis collinearity. Third, the lack of phenotype under optimal, unstressed laboratory conditions does not preclude a critical requirement for this subregion in the wild, where subtle variations in regulatory efficiency may be exposed to continuous environmental or physiological selective pressures that our assays do not capture. Further, our quantitative expression analysis was performed on whole embryos, so a change confined to a small domain of the fin bud, or a subset of neural tissue where CR2 drives reporter expression, would be diluted against unaffected tissue and could account for a real effect appearing only as a non−significant trend; tissue−dissected or single−cell approaches would be better suited to resolving this.

We also observed strand specific transcription from CRs during early embryonic development in a direction similar to *hoxd* genes and opposite to *evx2*. The expression profiles of both regions across developmental stages are also similar. Similarity in the spatio−temporal expression of these RNAs raises the possibility that a single, continuous long non−coding RNA (lncRNA) encompasses both elements. The functional requirement of these RNAs from CRs in *hoxd* regulation requires deeper investigation. RNAs from T−UCEs are known to have a variety of functions, including regulating gene expression via miRNAs, regulating splicing, binding to target genomic regions, and recruiting regulatory factors (Cajigas et al., 2018; Karginov et al., 2023; Lareau et al., 2007; Olivieri et al., 2016). The regulatory function of RNA from CRs would be worth further study. The unsolved questions in the field are: what is the biological significance of ultraconservation, and what biological mechanisms would require such a high level of sequence constraints? We hypothesize that these transcripts leverage sequence− specific base−pair complementarity to tether back to their own genomic loci. This structural feedback loop could facilitate recruitment of regulatory complexes, thereby tuning the transcriptional output of the adjacent *hoxd* complex. This self−tethering mechanism is speculative and, beyond the outline proposed here, remains entirely untested.

## Conclusions and future perspectives

Hox genes are known to be expressed beyond embryonic development and have non− homeotic functions as well. Since we have studied the role of CR2 in *hoxd* genes regulation during early embryonic development, it would be interesting to study the function of CR2 in the regulation of *hoxd* genes in adult tissues. Our results suggest strand−specific transcription spanning CR2 and CR3. Prior to gastrulation, RNA is ubiquitously expressed throughout the embryo; following the onset of gastrulation, its expression becomes restricted to the anterior region. Exploring the sequence features and functional roles of these CR−derived RNAs, including the possibility that they bind to their own genomic loci, could provide deeper insight into the evolutionary significance of their ultraconserved nature.

## Acknowledgements

We thank all present and former members of the RKM laboratory for helpful discussions and critical input. We also thank Dr. Megha Kumar for her suggestions and technical inputs throughout the study. We thank the CSIR−CCMB fish facility and the central instrumentation facility for their support.

## Author’s contribution

SY developed the concepts, designed and performed experiments, analyzed data and wrote the manuscript. AK performed genotyping, ISH and embryo collections. RKM conceived the project, developed the concepts, analyzed data and supervised the manuscript writing.

## Materials and methods

### Zebrafish husbandry and ethics statement

Zebrafish (*Danio rerio*) of the Tubingen background were maintained at 28.5 °C on a 14 h light / 10 h dark cycle under standard conditions. Embryos were obtained by natural spawning and staged according to Kimmel et al. (1995). All animal procedures were approved by the Institutional Animal Ethics Committee of CSIR−CCMB and were performed in accordance with the guidelines.

### Guide RNA design and cloning

For creating genetically modified fish lines, we used the CRISPR−Cas9 technique (Irion et al., 2014). Online available tools such as CRISPOR (Concordet & Haeussler, 2018), CHOP−CHOP (Montague et al., 2014) and E−CRISP (Heigwer et al., 2014) were used to design guide RNA (gRNA). Query sequences were provided for the region of interest. The reference genome used was *Danio rerio* (zebrafish), UCSC 2017 (GRCz11/danRer11). The protospacer adjacent motif sequence 5’−NGG−3’ associated with *Streptococcus pyogenes* was used to get target guides. gRNAs that were suggested in more than 2 tools, with minimum off−targets and higher specificity scores, were chosen for the experiments. Target guide forward and reverse oligos were purchased with BsaI competent overhangs (Supplementary Table 5). The equimolar guide oligos were then annealed in annealing buffer (10mM Tris pH 8, 50mM NaCl, 1mM EDTA) by first heating at 95°C for 2 min, followed by reducing the temperature to 85°C by a decrement of 2°C/sec and finally to 25°C by reducing 0.1°C/sec. The annealed double− stranded oligos were diluted 100 times and ligated using T4 DNA ligase into the BsaI−digested pDR274 plasmid. BsaI digestion site flanks the T7 promoter and the guide RNA scaffold in the plasmid. The ligated plasmid was transformed into DH5a competent cells. The positively transformed colonies were selected by growing LB agar plates containing kanamycin. Positive colonies were sequence−verified and then plasmid isolation was performed using Qiagen’s midi−prep kit following the user’s guide.

### Guide RNA synthesis

Plasmids with guide oligos inserted were linearised with HindIII, which digests downstream of the guide scaffold. In−vitro transcription reaction (IVT) (Invitrogen MEGAscriptTM T7 transcription kit, cat. No. AM1334) was set up to synthesise gRNA using the T7 promoter upstream of the guide insert. The transcribed RNA was purified using mini−Quick Spin Columns (Roche, cat. no. 11814427001) according to the manufacturer’s user protocol. Linearised plasmid, IVT product without DNase treatment, and with DNase treatment were run on 2% agarose gel to check the quality of gRNA. Purified gRNAs were quantified using a spectrophotometer. gRNAs were aliquoted into 2 μL in each PCR tube and stored at −80°C. For each injection set, fresh aliquots were used to avoid gRNA degradation. For all experiments, 150 pg of each gRNA and 350 pg of Cas9 enzyme, in a 3 nl volume, were injected into each embryo.

### Fin clipping and genomic DNA isolation

For genotyping, around one−month−old fish were anaesthetised using tricaine (150μg/ml). The caudal fin was amputated using a clean surgical blade and incubated in DNA extraction buffer (Tris pH 8.5−10mM, EDTA− 1mM, NaCl− 25mM, and Proteinase K− 200μg/ml) for at least 4h at 56°C. ProteinaseK was heat−inactivated at 95°C for 10min. Samples were centrifuged and stored at 4°C for short−term storage. For genetic screening, genomic DNA extracted from the clipped fins was used as a template. Two primers flanking the target sites of genetic manipulation were used for PCR−based screening for deletions. The primers used for the screening experiments are listed in the Supplementary Table 6. Fish that were positive for the desired genotype (founder fish, F0) were segregated and raised separately.

### Zebrafish line generation and maintenance

The founder fish was bred with wild−type (WT) fish, and the progeny were screened for the desired heterozygous genotype. Those heterozygous fish were then self−crossed to get the population of embryos with mixed genotypes. Those fish were genotyped at one month of age and segregated into different genotypes (WT, het, and homo). Parents were screened at regular intervals to prevent the mixing of genotypes. Sequence of different CR deletion alleles is given in Supplementary File 1 and genomic coordinates of the sequences in Supplementary Table 7.

### Probe synthesis for RNA-ISH

We used in vitro transcription to generate Digoxigenin labelled RNA probes for the target genes (Thisse & Thisse, 2014). The probe lengths used in the experiments ranged from 300 to 800 bp. Fragment of target genes (300−1000 bp) amplified using gene−specific forward and reverse primers with T7 (5’ TAATACGACTCACTATAGGG 3’) and SP6 (5’ ATTTAGGTGACACTATAGAA 3’) promoter overhangs at their 5’ end (Supplementary Table 8). Amplified DNA products were gel−purified and sequence−confirmed before using them in the downstream reaction. DNA templates (800ng−1000ng/ reaction) were used for one *in vitro* transcription (IVT) reaction to make sense/antisense probe. The DIG labeling kit (Roche, cat. No. 11175025910) was used to prepare a digoxigenin−labeled RNA probe according to the manufacturer’s protocol. The Promotor (T7/SP6) for transcription was chosen based on the required probe (anti−sense/sense) for IVT. After RNA synthesis and DNase treatment, probes were purified using mini−Quick Spin Columns (Roche, cat. No. 11814427001). The integrity of the RNA probe and the presence of DNA template contamination were checked by running the samples on a 2% agarose gel. For the first ISH, both sense and antisense probes were used to assess probe specificity. For each ISH experiment, we have used a 0.5 ng/ul probe concentration diluted in hybridization mix buffer.

### PTU treatment

Retinal pigmentation begins after the 24−hour post−fertilisation (24hpf) stage, which later spreads along the body axis. It is challenging to visualize any kind of staining signal in pigmented embryos. To avoid such issues, embryos were raised in 0.003% PTU in embryo media in the dark from 24 hpf onwards.

### RNA-*in situ* hybridization (RNA-ISH)

For RNA−ISH, we followed the standard protocol for probe synthesis, embryo preparation, probe hybridization, and staining (Thisse & Thisse, 2014). Embryos were collected and fixed in 4% paraformaldehyde (PFA) overnight at 4°C. The next day, the embryos were sequentially dehydrated in methanol (25%, 50%, 75%, and 100%) and stored at −30°C until further use. For each experiment, at least 25 embryos were collected per sample, and the experiment was repeated at least 3 times. Methanol−fixed samples were rehydrated in a series of PBST solutions (25%, 50%, 75%, and 100%). Embryos were washed twice with PBST for 10 min at room temperature. Embryos were washed with pre−warmed (at 65°C) hybridization wash buffer (HW). After that, embryos were incubated with a hybridization mix (HM) at 65°C for 4h. HM was replaced with pre−warmed, labeled RNA probes diluted in HM (0.5 ng/μl) and incubated at 65°C overnight. Next day, the probes were collected for reuse. Embryos were washed with HM, 75% HW+25% 2XSSC, 50% HW + 50% 2XSSC, 25% HW + 75% 2XSSC and 2XSSC for 10 min each at 65°C followed by 0.2XSSC twice, 20min each at 65°C. Samples were then transferred to RT and successively transferred to 1XTBST. Blocking was done with a blocking buffer (10% heat−inactivated fetal bovine serum + 1X TBST) for 2h at RT. Samples were incubated overnight at 4 °C on a shaker with the anti−DIG antibody conjugated to alkaline phosphatase (Roche, cat. No. 11093274910), diluted 5000−fold in blocking buffer. The next day, the antibody solution was removed, and the samples were washed 4 times with 1XTBST, 10min each at RT. Samples were then washed with an alkaline buffer (NTMT) 4 times, 10 min each at RT. An equal volume of NTMT and 1−Step NBT−BCIP substrate solution (Thermo Scientific, cat. no. 34042) was mixed and added to the embryo samples. Embryos were periodically monitored to check the signal development. Once the proper signal was achieved, the reaction was stopped by replacing the substrate−containing solution with 4% PFA and incubating at RT for 20 min. After that, samples were washed 3 times with 1XPBST at RT and stored in PBS until imaging. In experiments comparing the expression of target genes in WT and genetically modified embryos, ISH reactions were stopped simultaneously.

### Skeletal and cartilage double-staining

Skeletal and cartilage double staining was performed following the protocol described by Walker and Kimmel, 2007 (Walker & Kimmel, 2007). Larvae of desired stages were fixed in 4% paraformaldehyde at 4°C overnight. The next day, samples were washed twice with PBST. Samples were dehydrated in 50% ethanol for 10 min at RT. Freshly prepared acid−free double− staining solution 10μl of solution B in 1ml of solution A (solution A− 0.03% alcian blue, 125mM MgCl2 and 70% ethanol, and Solution B− 0.5% alizarin red) was added to the samples and left on shaking at RT overnight. After removing the staining solution, the samples were washed with water for 5 minutes. Then, the bleaching solution (1.5% H2O2 and 1% KOH) was added, and the samples were incubated in the dark with the lids open until the eye and body pigments were cleared (approximately 30 minutes). Then, tissue clearing was done by adding 20% glycerol and 0.25% KOH at RT for 2h to overnight. After tissue clearing, 50% glycerol and 0.1% KOH solution were added to the samples, and the fish were imaged the same week.

### RNA isolation, cDNA preparation and RT-qPCR

Zebrafish embryos/tissues were fixed in TRIzol and stored at −80°C until the next use. RNA was prepared using a Macherey−Nagel (MN) NucleoSpin RNA kit following the manufacturer’s protocol. RNA integrity was checked by running 500 ng of RNA on an agarose gel. For cDNA preparation, 2 μg of RNA was used in a 20 μl reaction. Oligo dT primers were used to make a cDNA library using the SuperScriptTM IV first−strand synthesis system by Thermo Fisher Scientific (cat. No. 18091050), following the user’s guide protocol. 1−5ng cDNA was used for 10μl RT−qPCR in 384−well PCR plates. PowerUp SYBR Green Master Mix from Applied Biosystems was used to set up PCR using Quant Studio VII software, following the protocol suggested for the aforementioned enzyme master mix. The primers used for RT−qPCR are listed in Supplementary Table 9.

### Statistical analysis

All analyses were performed in GraphPad Prism 8.0.2. Relative expression of *hoxd* complex genes was assessed on ΔCt values using unpaired multiple t−tests, with four independent embryo pools per genotype at each stage. The p−values in Supplementary Table 4 are uncorrected for multiple comparisons and should be read as exploratory. Observed genotype counts were compared with the expected 1:2:1 ratio using the chi−square test. Somite counts were compared using unpaired two−tailed t−test. Values are reported as mean ± SEM unless stated otherwise.

