## Supplementary Figure/Table/File1 for "Functional dissection of ultraconserved non-coding regions present near the *Hoxd* complex in vertebrates"

**[2] Academy of Scientific and Innovative Research (AcSIR), Ghaziabad, Uttar Pradesh, 201002, India**

**[3] Tata Institute for Genetics and Society (TIGS), Bengaluru, Karnataka, 560065, India**

**Supplementary Figures**

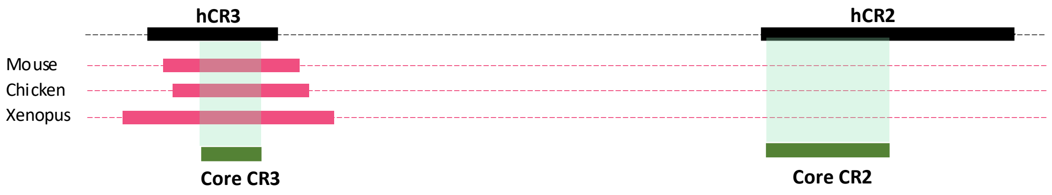


**Supplementary figure 1: ESTs mapping to CRs loci.** Schematic representation of ESTs from mouse, chicken and xenopus overlapping with CR3. Pink blocks represent ESTs.

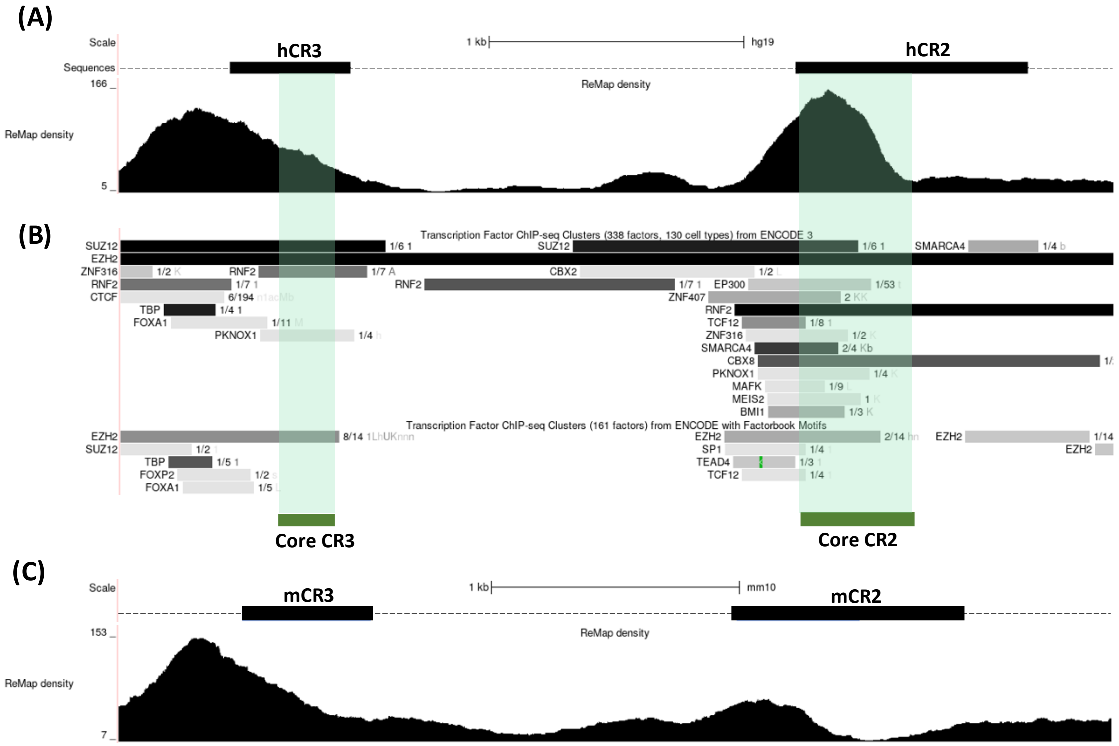


**Supplementary figure 2: Regulatory factors binding to the CRs loci.** (A) ReMap density track comprises integrative density of Transcription factor binding (scale 5-166) obtained by integrating all ChIP-seq data from GEO, ENCODE and ArrayExpress for all transcription regulators in different human cell types and tissues. (B) The transcription factor ChIP-Seq cluster track displays experimentally validated TF binding regions from two projects. The top set is from ENCODE project, where 338 transcription factors were tested across 130 cell types. The bottom set represents 161 factors from the ENCODE project, along with binding motifs identified by the Factorbook repository. In this track, the grey box represents the peak cluster of transcription binding, and its intensity reflects the highest signal strength observed across all cell lines contributing to the peak cluster. The left side of the grey box has the name of the transcription factor, and the right side has details about how many cell types (with single-letter abbreviations representing the cell types) have a binding signal for the transcription factor from the total cell types assessed for the mentioned transcription factor. Single letter abbrevations for cell types: 1- H1-hESC, a-A673, b- bipolar neuron, c-cardiac muscle cell, M- MCF-7, n- Neural progenitor cell, K-K562, t- tibial nerve, A- A549, L-HepG2, h- HEK293T (C) ReMap density track comprises integrative density of Transcription factor binding (scale 7-153) obtained by integrating all ChIP-seq data from GEO, ENCODE, and ArrayExpress for all transcription regulators in different mouse cell types and tissues.

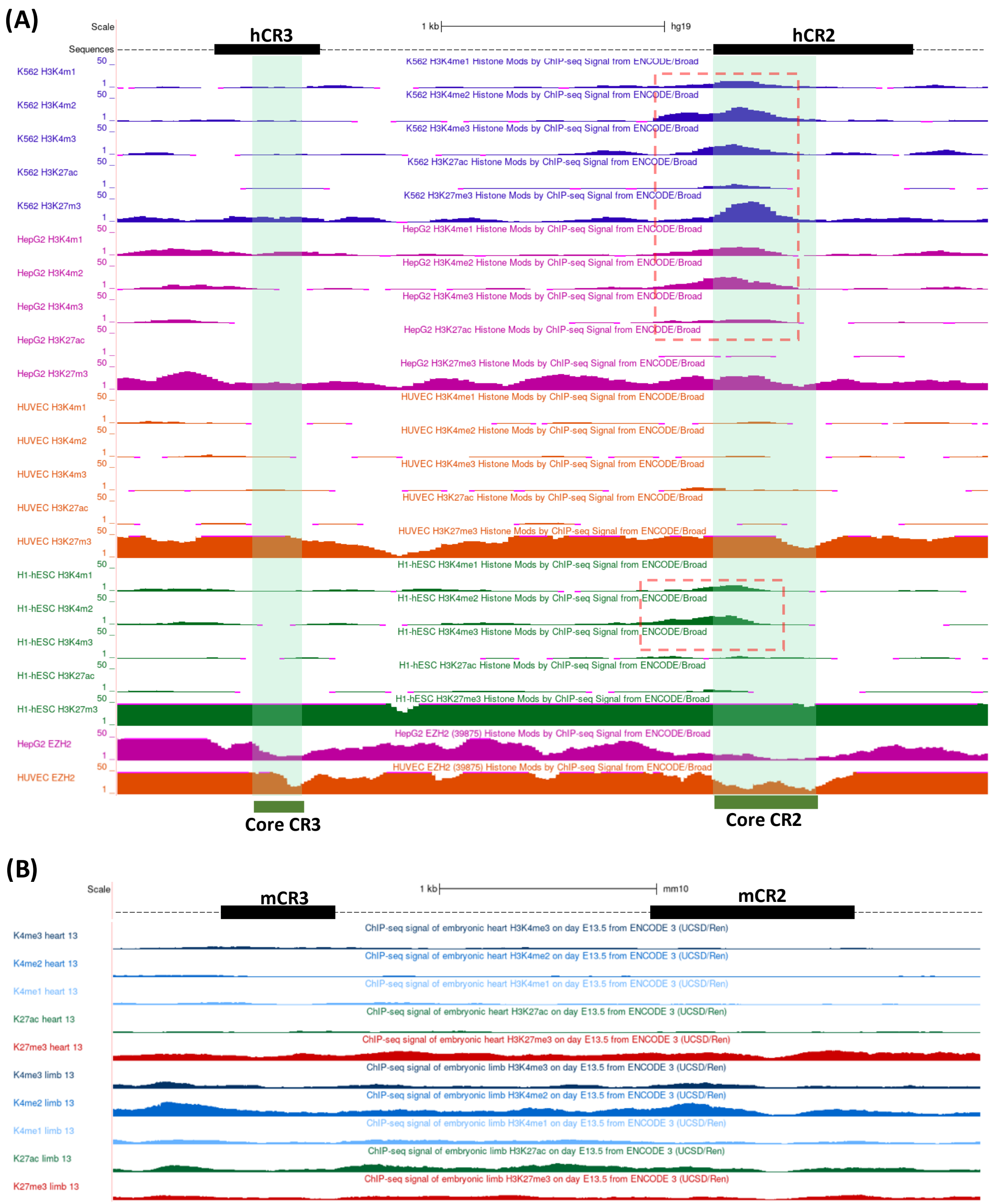


**Supplementary figure 3: Histone modifications in CRs loci.** (A) The track maps the ChIP sequence signal of various histone modifications over the CR locus in different human cell types generated by the Broad ENCODE group. EZH2 protein binding in HepG2 and HUVEC cell lines is also represented at the bottom. (B) The track maps the ChIP signal for various histone modifications across the CR locus in limb and heart tissues of the E13.5 mouse embryo, as part of the ENCODE project.

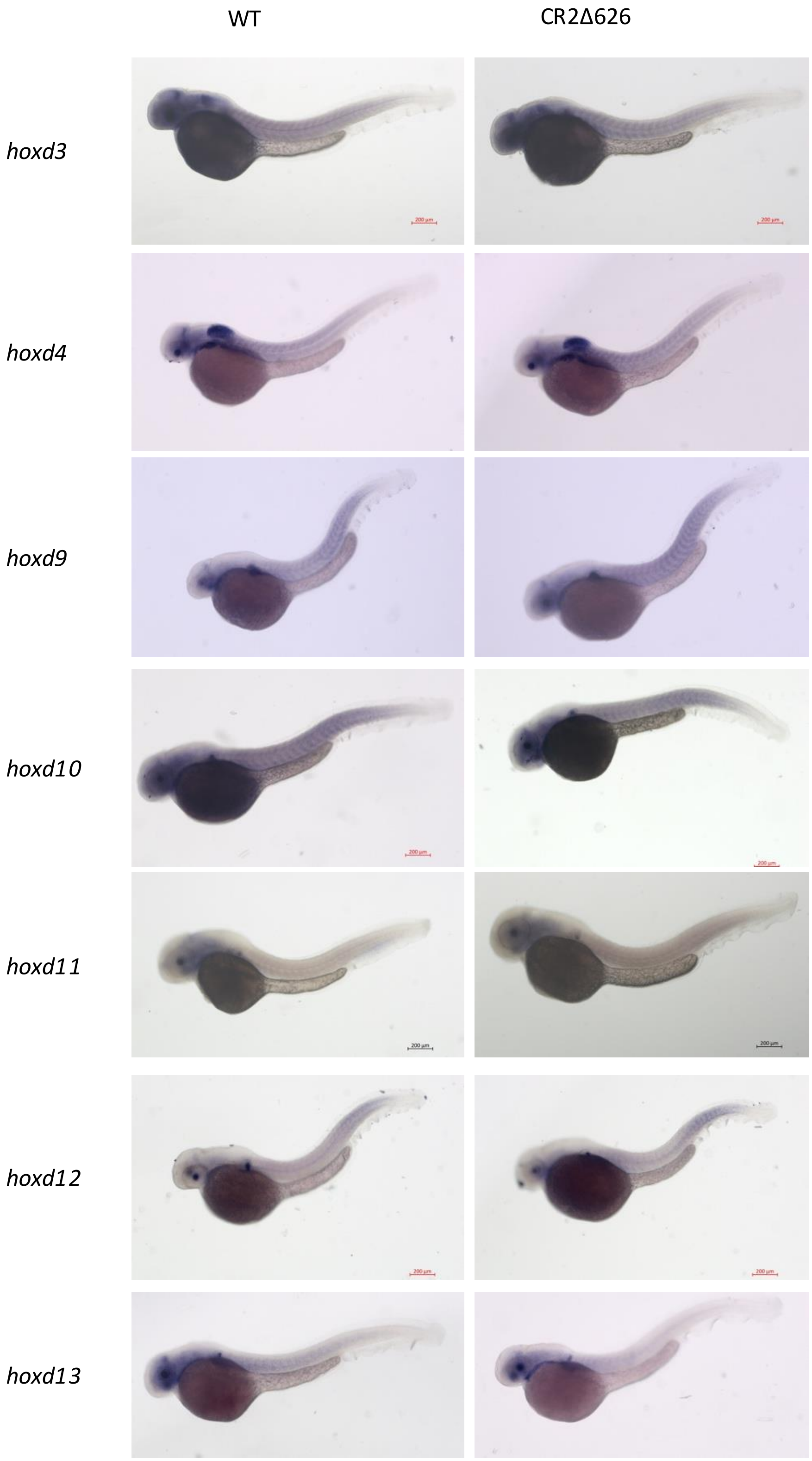


**Supplementary figure 4: Expression of *hoxd* complex genes in 48 hpf CR2Δ626 embryos.** RNA-ISH of all the *hoxd* complex genes and *evx2* gene in 48 hpf WT and CR2Δ626 embryos.

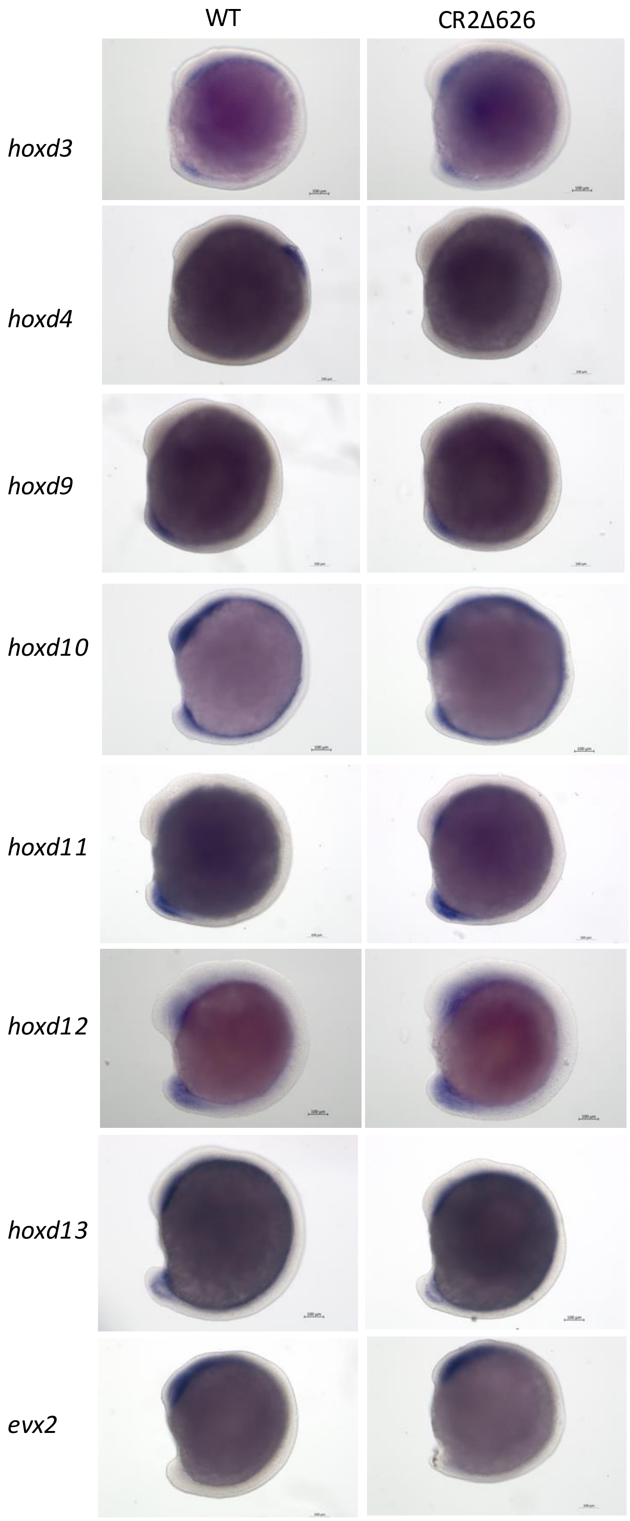


**Supplementary figure 5: Expression of *hoxd* complex genes in 16 hpf CR2Δ626 embryos**
RNA-ISH of all the *hoxd* complex genes and *evx2* gene in 16 hpf WT and CR2Δ626 embryos.

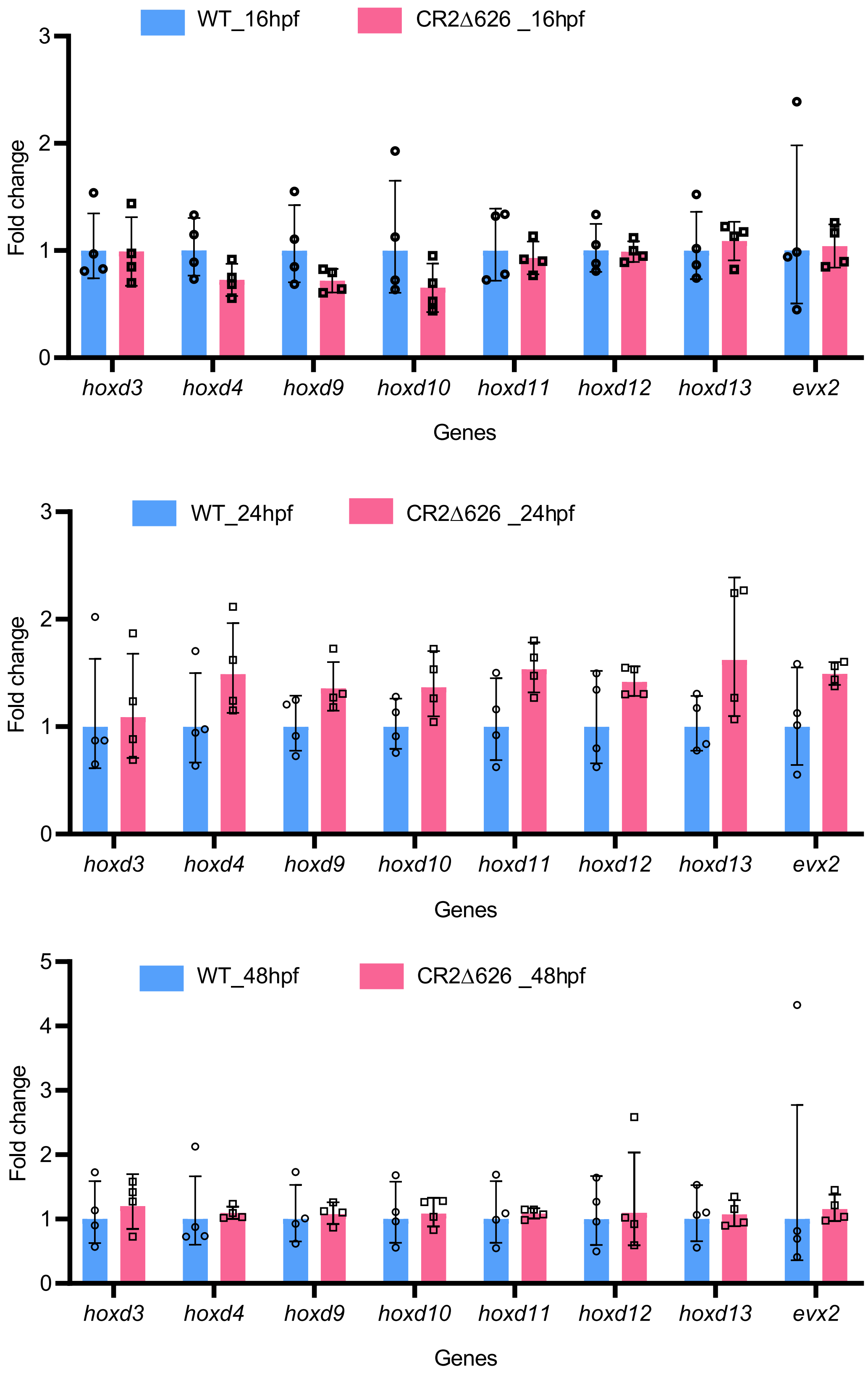


**Supplementary figure 6: Relative expression of all the *hoxd* genes and *evx2* gene.** Relative expression of hoxd and evx2 genes in 16 hpf (A), 24hpf (B) and 48hpf wild type vs CR2Δ626 embryos.

**Supplementary Tables**

**Supplementary Table 1. Percentage sequence conservation of CRs in different vertebrate species**

| Human | Chimpanzee | Mouse | Chicken | Xenopus | Zebrafish |
|---|---|---|---|---|---|
| Extended CR1 (400bp) | 99.25 | 97.79 | 94.51 | 78.66 | 72.30 |
| Core CR1 (200bp) | 100.00 | 99.00 | 97.01 | 90.96 | 79.49 |
| Extended CR2 (900bp) | 99.66 | 96.31 | 95.69 | 85.07 | 87.62 |
| Core CR2 (450bp) | 99.78 | 99.50 | 97.34 | 95.68 | 94.27 |
| Extended CR3 (550bp) | 99.00 | 98.71 | 96.61 | 86.59 | 88.75 |
| Core CR3 (200bp) | 99.50 | 99.50 | 99.50 | 95.43 | 92.54 |

**Supplementary Table 2. Summary table of CR deletion experiment**

| gRNA used | Expected size of deletion | No. of embryos injected | No. of fish screened | No. of founder fish | Germline transmission |
|---|---|---|---|---|---|
| CR2_gRNA1/ CR2_gRNA2+ CR3_gRNA3 | Around 2090bp | 2000 | 200 | 0 | NA |
| CR2_gRNA1/ CR2_gRNA2 + CR3_gRNA1 | Around 1900bp | 2000 | 200 | 1 | No |
| CR3_gRNA1 + CR3_gRNA3 | Around 430bp | 3000 | 1000 | 0 | NA |
| CR3_gRNA2 + CR3_gRNA3 | Around 230bp | 1500 | 800 | 0 | NA |
| CR2_gRNA1/2 + CR2_gRNA5 | Around 1100bp | 2000 | 1000 | 1 | No |
| CR2_gRNA1 + CR2_gRNA4 | Around 626bp | 2000 | 1000 | 3 | Yes |

**Supplementary Table 3. Genotype distribution of embryos obtained for CR2Δ626 heterozygous X heterozygous fish. Chi-square test for the expected 1:2:1 ratio.**

| Development stage | wt | het | homo | p-value |
|---|---|---|---|---|
| 2dpf | 17 (23.6%) | 38 (52.7%) | 17 (23.6%) | 0.8948 |
| 15dpf | 19 (29.7%) | 28 (43.75%) | 17 (26.5%) | 0.5698 |

**Supplementary Table 4: Statistical comparison of *hoxd* complex gene expression between wild-type and CR2 p.del embryos at 16, 24 and 48 hpf**

| Genes | p-value_16hpf | p-value_24hpf | p-value_48hpf |
|---|---|---|---|
| *hoxd3* | 0.835093 | 0.800535 | 0.982341 |
| *hoxd4* | 0.093508 | 0.156387 | 0.245409 |
| *hoxd9* | 0.126906 | 0.091436 | 0.218948 |
| *hoxd10* | 0.17178 | 0.099079 | 0.297638 |
| *hoxd11* | 0.671051 | 0.077593 | 0.230398 |
| *hoxd12* | 0.912916 | 0.154993 | 0.403283 |
| *hoxd13* | 0.696059 | 0.081917 | 0.319785 |
| *evx2* | 0.941056 | 0.121809 | 0.303326 |

**Supplementary Table 5: List of guide oligos used for CRISPR-cas9 deletion of CRs**

| Target gRNA | Sequence |
|---|---|
| CR2_gRNA1 | GGAAAATGTGTTAACAGAGC |
| CR2_gRNA2 | GAGGACAAAATCCGAACTAC |
| CR2_gRNA3 | GGGCGTGATTGGACAGTAGC |
| CR2_gRNA4 | TTGTTGTACGGGTTGTAACC |
| CR2_gRNA5 | GTAGTGTACAGCAGAGCTCC |
| CR3_gRNA1 | GTTCGTGGTAACACAAGCAC |
| CR3_gRNA2 | TAGTAATTCAGATACAAGCT |
| CR3_gRNA3 | GTCTAGTTTGCAGGAGAAAATG |

**Supplementary Table 6: Primers used for genotypic screening of CR deletions**

| Primer name | Sequence |
|---|---|
| CR2_screen_FP | GAGTAAATTCTGAATTGCAAAC |
| CR2_screen_RP | GTTTCATCTGATATAATTCTCC |
| CR3_screen_FP | ACACCGAGTCAATGATCGCC |
| CR3_screen_RP | AGGTGCAAATGTTCGACAGC |

**Supplementary Table 7: Genomic coordinates of CRs deletion alleles**

| Deletion allele | Deletion (GRCz13ab, chr9) | Size |
|---|---|---|
| CR2 complete | NC_141029.1:g.2192546_2193656del | 1111bp |
| CR2Δ626 | NC_141029.1:g.2192656_2193281del | 626bp |
| CR2+ region between CR2 and CR3 | NC_141029.1:g.2194506_2192608del | 1899bp |

**Supplementary Table 8: List of primers used for RNA-ISH**

| Primers name | Sequence (small letters are overhang) |
|---|---|
| Hoxd3_FP_T7 | taatacgactcactatagggTTCTTTGGAGAACCCGGCAA |
| Hoxd3_RP_SP6 | atttaggtgacactatagaaGTCGGGGGAAACTTCGTTGA |
| Hoxd4_FP_T7 | taatacgactcactatagggAAAACAGCAGAATAGGGATGG |
| Hoxd4_RP_SP6 | atttaggtgacactatagaaAGCCCAGGCTACTACAGTCC |
| Hoxd9_FP_T7 | taatacgactcactatagggCGGTCCTTTTCGGGGGTAAA |
| Hoxd9_RP_SP6 | atttaggtgacactatagaaTGTTCGGAGGCAAGGTTTCG |
| Hoxd10_FP_T7 | taatacgactcactatagggATTCTTCTCCTCCAGCGTGC |
| Hoxd10_RP_SP6 | atttaggtgacactatagaaACATGGACATGACCGTCCAC |
| Hoxd11_FP_T7 | taatacgactcactatagggTTCTCGGAGTTTGCCGTCTC |
| Hoxd11_RP_SP6 | atttaggtgacactatagaaTTCTCGTCGGTTTCGTCGTT |
| Hoxd12_FP_T7 | taatacgactcactatagggCCATCGGTGAAAGCTGGTCT |
| Hoxd12_RP_SP6 | atttaggtgacactatagaaGGCTATGTCGCTCCCCTTTT |
| Hoxd13_FP_T7 | taatacgactcactatagggGGTTTCCAGATATGCGGGCT |
| Hoxd13_RP_SP6 | atttaggtgacactatagaaCAAAGTTCCACAGAACGCCG |
| evx2_FP_T7 | taatacgactcactatagggGGATATACGCGTCTGGCCGAT |
| evx2_RP_SP6 | atttaggtgacactatagaaTCGCTGTTCATGTCGGAGTC |
| Xirp2a_FP_T7 | taatacgactcactatagggGACTTCCTGTTGCAGAGCAAG |
| Xirp2a_RP_SP6 | atttaggtgacactatagaaGGATGCTCTGTTGCTGAACT |

**Supplementary Table 9: List of primers used for RT-PCR**

| Gene name | Forward primer | Reverse primer |
|---|---|---|
| *hoxd3* | GTACGGTCCAACACACCAGG | TCGCAGGTTTCTCCTGCTGCTGGGC |
| *hoxd4* | GTGCACGTTACTACGGTGAACCCG | TTCATTCTCCTGTTCTGAAAC |
| *hoxd9* | ACAACAACTTGACCCAAGCAAC | TCACCTGGCGCTCTGTTAAA |
| *hoxd10* | CACACAGAGAAGGCAAAGAATCTA | CTGTCAGTCAGGTTGACGCT |
| *hoxd11* | AGACGGCAAACTCCGAGAAG | ACTTTGTCGCACTCGAGCC |
| *hoxd12* | CATAGACACCAATGGACTGCC | TCATCCTGCGATTCTGAAAC |
| *hoxd13* | AAACCATCATTAACAGAGGAAGC | TAGAAGAAGCGATCCGCCG |
| *evx2* | TTTGGGAACCACCACAACGA | TTGCTGTCCGAGAAACCTTTA |
| *gapdh* | GTGGAGTCTACTGGTGTCTTC | GTGCAGGAGGCATTGCTTACA |

**Supplementary File 1**

Sequence of 626bp subregion deletion in CR2:

TAGCGGGAGGATCATCGATCTTTGCATTTCTATCTTGCTGGTGGACATTTAATTTGGGTTTTCTATTAGCACCGAA
ATAAAGAAAAAAATAAAATATATTAAACAACCCGCAGATTTATTTTCTTGTCAGAAGCAATTCGCCTAAGGTGACA
GACAGTTTGCTGCATTTTATGATGAATTCTTTTTTCAGTCTTTGCACATTGGATACAAAACTGATGGCGTTATTGTG
GCCAGTGTTTTTCTGCCCTCTGTCTTCTCCAGTTTGCCTCTCCATCTATCTCAATGCCTTTGTTGTACGGGTTGTAA
CCCTCGCCTTGGTAAAGAGCATAACGAAACATGGCGATACATCACACTCTTGCTGCGCCTCTAATCCCATTTCTTT
AATGGGGACGTTCAAAAAGAAAGCAACTTATCTCAAAGTCCCCCGGGAGTGTGCCGGTATATATTTAACCAGGC
TGCAATTAACGACAGTATCAGCTTGTATCATTTAGAGGTAATGTAAAAATAATGGTAAATTATAGGGCCGGAATGA
CCCGTGATCGCAGACTCCATATTCCCCATAGTTTTAGTGTCGTTTTAATTAATGCCAGACTCACTTTCTCACTAAGG
AGAAAGGGAAAATGTGT

Sequence of entire 1111bp CR2 deletion:

GCCCCACGTGCTCCCTGTAGTTCGGATTTTGTCCTCCCCGCAGCAGCTGTCACCTCGCATTACTCGCAGTCAGCTA
AATGAAACATTATTCTAAACATATGCATCGTAATCAGTTCGGTCACACTTACAAGAACACGCGTTAATAAGGCAAT
CAATCACAGTGGAACAAGCGAGTTGTTCTGCAACAGCTCATAAACAGTGTGTAATGAAGAATTGGAGGTTAGCA
TGACACGCGCTGATCAGATAATCAATGTCAAAGATGCGATGAATGTCAGTAAATGTAGTTTTTATGTCGTTTCTATA
AAATCTCCAATTTACAACAAGCAGTTCAATTACCCAGAAAATACAGTCAATTAAACGGGCGTGATTGGACAGTAG
CGGGAGGATCATCGATCTTTGCATTTCTATCTTGCTGGTGGACATTTAATTTGGGTTTTCTATTAGCACCGAAATAA
AGAAAAAAATAAAATATATTAAACAACCCGCAGATTTATTTTCTTGTCAGAAGCAATTCGCCTAAGGTGACAGAC
AGTTTGCTGCATTTTATGATGAATTCTTTTTTCAGTCTTTGCACATTGGATACAAAACTGATGGCGTTATTGTGGCC
AGTGTTTTTCTGCCCTCTGTCTTCTCCAGTTTGCCTCTCCATCTATCTCAATGCCTTTGTTGTACGGGTTGTAACCCT
CGCCTTGGTAAAGAGCATAACGAAACATGGCGATACATCACACTCTTGCTGCGCCTCTAATCCCATTTCTTTAATG
GGGACGTTCAAAAAGAAAGCAACTTATCTCAAAGTCCCCCGGGAGTGTGCCGGTATATATTTAACCAGGCTGCA
ATTAACGACAGTATCAGCTTGTATCATTTAGAGGTAATGTAAAAATAATGGTAAATTATAGGGCCGGAATGACCCG
TGATCGCAGACTCCATATTCCCCATAGTTTTAGTGTCGTTTTAATTAATGCCAGACTCACTTTCTCACTAAGGAGAA
AGGGAAAATGTGTTAACAGAGCAGGAGCTCTGCTGTACACTACTGGATACATTTTTTTATCGTTTAAAAAAATATT
TTAAAAGGAAAAATTTTAAATACTTAATTTAAAAATCACATTTTTAA

Sequence of 1899bp deletion spanning CR2 and the region between CR2 and CR3:

CTTTGTTGAGGACTGTTTAACGGGCGATCATTGACTCGGTGTTATCAGCTTATCAGTCGTTGTGTAGGTGTATAAA
TGAAGACTTAGGCTATGGTTAGACGTGAATTAGATTTTCACTGACAGGCCAAATTTAGCCTGGTTTTAGCCTAGA
TCAAAATTCACCGGCTATTATACGGATATATTCGTCTATTAAGACCTGTCTAATATTAGGCTATGGTTAGACGTCTAT
CTTCGCTGACACCCCAAATTAAGCCTAGGCTATGTTTAGACGGCTAATAGACTTTATATATATATATATATATATATA
TATATATGCTTTCTGGGTAGTAGATGCGCGTTGTGTTTACTTATTTAAATCACGTTTCATCTGATATAATTCTCCATAT
TACCAAAAGACCATGTTTTTTTTTATCATTTTTTGGATGCAATTCATTTACAGGAAAATGCATGATAACATATGGCG
AGGTAAACCCTATAATATCACCCCCCAGCATAAAAGACTTCATTTATCGTTAGCACGTCGAGCGTTTCAGTTCAATC
GCAGCAACTGGATGGGATCACCCTCACCCCTTTCAGACCACCTGAAAATAATACAACCTGACAGCTTTGGGTGG
GCTTTACCAAATTATTCTATTACGTATTTGATTAAAAACTGTCTGATTGGCAGCGTTGACTCTCCACGGCACAATGC
GTTATATAGTTCTCCAAAAAATGTAGCGCGCAAGTTCAGTCGCAGGGCATCGCTTTGCGCCCGCGATTCGTATTT
GTGAAGTTCCACCTTAAAACAACAGTGTTTTGGCCCCACGTGCTCCCTGTAGTTCGGATTTTGTCCTCCCCGCAG
CAGCTGTCACCTCGCATTACTCGCAGTCAGCTAAATGAAACATTATTCTAAACATATGCATCGTAATCAGTTCGGTC
ACACTTACAAGAACACGCGTTAATAAGGCAATCAATCACAGTGGAACAAGCGAGTTGTTCTGCAACAGCTCATA
AACAGTGTGTAATGAAGAATTGGAGGTTAGCATGACACGCGCTGATCAGATAATCAATGTCAAAGATGCGATGA
ATGTCAGTAAATGTAGTTTTTATGTCGTTTCTATAAAATCTCCAATTTACAACAAGCAGTTCAATTACCCAGAAAAT

ACAGTCAATTAAACGGGCGTGATTGGACAGTAGCGGGAGGATCATCGATCTTTGCATTTCTATCTTGCTGGTGGA
CATTTAATTTGGGTTTTCTATTAGCACCGAAATAAAGAAAAAAATAAAATATATTAAACAACCCGCAGATTTATTTT
CTTGTCAGAAGCAATTCGCCTAAGGTGACAGACAGTTTGCTGCATTTTATGATGAATTCTTTTTTCAGTCTTTGCA
CATTGGATACAAAACTGATGGCGTTATTGTGGCCAGTGTTTTTCTGCCCTCTGTCTTCTCCAGTTTGCCTCTCCATC
TATCTCAATGCCTTTGTTGTACGGGTTGTAACCCTCGCCTTGGTAAAGAGCATAACGAAACATGGCGATACATCAC
ACTCTTGCTGCGCCTCTAATCCCATTTCTTTAATGGGGACGTTCAAAAAGAAAGCAACTTATCTCAAAGTCCCCCG
GGAGTGTGCCGGTATATATTTAACCAGGCTGCAATTAACGACAGTATCAGCTTGTATCATTTAGAGGTAATGTAAA
AATAATGGTAAATTATAGGGCCGGAATGACCCGTGATCGCAGACTCCATATTCCCCATAGTTTTAGTGTCGTTTTAA
TTAATGCCAGACTCACTTTCTCACTAAGGAGAAAGGGAAAATGTGTTAACAGAGCAGGAGCTCTGCTGTACACT
ACTGGATACATTTTTTTATC
